# A long-lived Human Neurovascular PENTA Culture Model Reveals Incomplete Vascular Repair and Glial-Mediated Signaling After Traumatic Brain Injury

**DOI:** 10.1101/2025.08.18.670951

**Authors:** Daniel S Hinrichsen, Sunghyun Jun, Charitha C Anamala, Mitchell D Thielen, Anapaula Garcia, Volha Liaudanskaya

**Affiliations:** Biomedical Engineering Department, University of Cincinnati, Cincinnati, USA; Neuroscience Graduate Program, University of Cincinnati, College of Medicine, Cincinnati, OH, USA

## Abstract

Traumatic brain injury (TBI) frequently leads to chronic neurovascular dysfunction, yet mechanistic insights into human-specific responses have been limited by the absence of long-term, multicellular in vitro models. Here, we report a five-cell-type human neurovascular culture system, comprising endothelial cells, astrocytes, pericytes, microglia, and neurons, engineered within a 3D scaffold to study injury-induced remodeling over multiple weeks. This PENTA-culture platform recapitulates hallmark features of the neurovascular unit and enables dissection of cell-specific contributions to vascular repair and degeneration. Upon mechanical trauma, cultures exhibit a biphasic response marked by acute endothelial disintegration, mitochondrial stress, and glial activation, followed by a delayed and incomplete repair. Confocal and proteomic analyses reveal persistent disruptions in tight junction organization, elevated TDP-43 and APP expression, and altered angiogenic and immunomodulatory signaling involving Tie2 and JAK/STAT pathways. Compared to simpler culture systems, the inclusion of microglia and neurons enhances post-injury cytokine resolution and junctional recovery, underscoring the importance of neuroimmune crosstalk. This system offers a mechanistically rich, human-relevant model for studying chronic neurovascular dysfunction and therapeutic revascularization.

## Introduction

Traumatic brain injury (TBI) affects more than 69 million individuals globally each year and remains a major cause of long-term neurological disability ^1–3^. While acute injury symptoms, such as edema, hemorrhage, and neuronal damage, often receive immediate clinical attention, it is the delayed secondary injury cascade that drives progressive cognitive decline, psychiatric disorders, and neurodegeneration ^3,4^. This cascade, which unfolds over days to months, involves blood–brain barrier (BBB) breakdown, chronic inflammation, glial activation, and impaired metabolic signaling ^5–7^. Repetitive mild TBIs, common in military personnel and athletes, further exacerbate these outcomes and are strongly linked to the development of chronic traumatic encephalopathy (CTE) and Alzheimer’s disease (AD)^8–11^. Vascular dysfunction is increasingly recognized as a central initiator and amplifier of these processes, underscoring the need for mechanistic insight into the cellular interfaces that regulate brain homeostasis after injury ^12–14^. The multifaceted nature of TBI, encompassing both immediate and delayed effects, underscores its significance as a public health challenge and a critical area of study in biomedical engineering.

At the core of this interface lies the neurovascular unit (NVU), an integrated assembly of endothelial cells, pericytes, astrocytes, neurons, and microglia that governs cerebral blood flow, BBB selectivity, and neuroimmune responses ^3,5,15–17^. The NVU dynamically adapts to local metabolic demand and neural activity, but is highly vulnerable to mechanical and inflammatory insults ^13,14,17,18^. Following TBI, NVU disruption compromises barrier integrity and intercellular communication, propagating neuroinflammatory and degenerative cascades ^5,15,18–20^. While existing preclinical models, particularly rodent systems, have provided foundational insights into these events, their translational relevance is limited by interspecies differences in brain structure and immune signaling ^9,20–23^. In vitro models, including transwell co-cultures and 3D endothelial scaffolds, offer valuable platforms for mechanistic studies and molecular screening, but often lack long-term stability, vascular fidelity, or the full complement of NVU cell types ^24–27^

Here, we present a modular, vascularized 3D human brain model incorporating all five principal NVU cell types (“PENTA” cultures) to dissect how vascular injury contributes to neurodegeneration following moderate TBI. Built on a silk-based scaffold optimized for extracellular matrix composition and cellular density, PENTA cultures exhibit sustained vessel-like architecture, tight junction protein expression, and injury-responsive cytokine secretion that recapitulate key aspects of human NVU physiology. By systematically isolating vascular and neuroimmune compartments and applying a controlled cortical impact protocol, we uncovered a biphasic injury response marked by early endothelial disintegration and mitochondrial stress, followed by incomplete structural recovery. Cytokine profiling revealed activation of chemokine-mediated signaling, JAK/STAT, and Tie2-associated angiogenic pathways— molecular signatures that mirror in vivo TBI pathology but have not previously been captured in a human in vitro NVU model. Importantly, inclusion of neurons and microglia shifted the secretome toward a pro-regenerative, immune-modulated profile, emphasizing the critical role of cellular complexity in vascular repair. This physiologically relevant, mechanistically rich platform provides an adaptable testbed for uncovering glial–vascular cross-talk and identifying therapeutic targets for TBI and related neurodegenerative diseases.

## Results

### Cellular density is critical in the formation of vessel-like structures

To address the overarching goal of engineering vascularized human brain tissue constructs, we first evaluated the role of endothelial cell density in promoting the formation of stable, vessel-like networks, via assessment of the ability of human brain microvascular endothelial cells (hBMECs) to self-organize into anatomically relevant/similar networks in the absence of other cell types.

As shown in Figure 1A, cultures with 0.5, 1, or 2 million GFP-expressing hBMECs were monitored weekly using epifluorescence microscopy to track the formation and persistence of the vascular network. At week 1, all three cell densities exhibited comparable network-like structures, characterized by branched morphologies and interconnected endothelial cords. However, as shown in Supplementary Figure 1A, by week 2, the lowest density group (0.5 million) displayed substantial regression of vascular-like organization, while 1 and 2 million cell culture densities maintained visibly intact networks. Confocal images, seen in Figure 1B, demonstrated elongated CD31+ and GFP+ tube-like structures present at 2 weeks after seeding. Shown in Supplementary Figure 1B, vascular network organization metrics (collected using AngioTool), including lacunarity, junction density, and average vessel length, did not differ significantly across cell seeding density conditions. By week 5, none of the groups retained long-term vascular structural integrity, indicating that endothelial cells alone are insufficient for chronic vascular development and maintenance, as shown in Figure 1A-B.

**Fig. 1:**
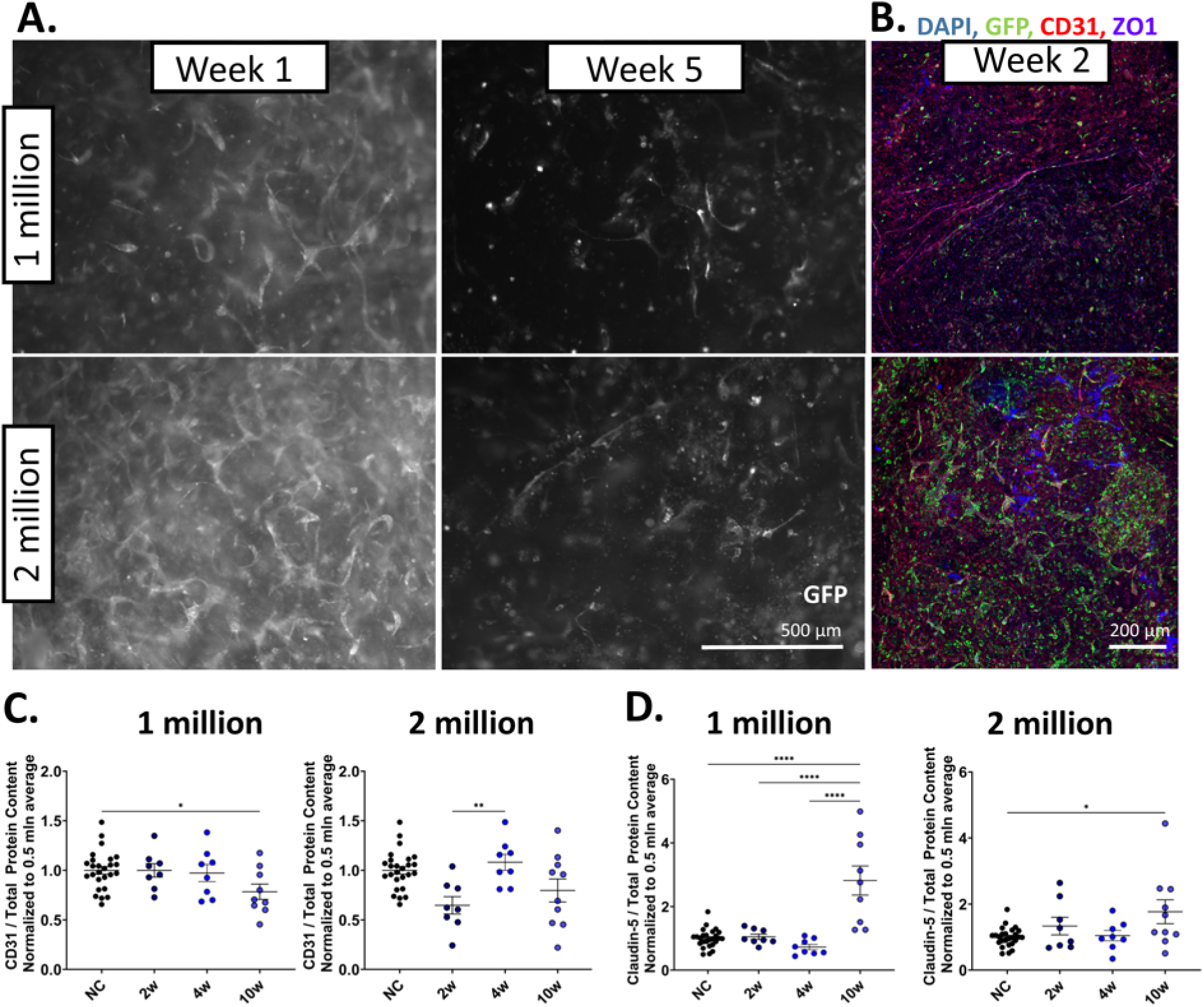
Cellular density regulates short-term vascular network formation but is insufficient for long-term maintenance in endothelial monocultures. GFP-labeled human brain microvascular endothelial cells (hBMECs) were seeded into collagen I–embedded silk scaffolds at three initial densities (0.5, 1, or 2 million cells) and maintained in culture for up to 5 weeks (W1–W5). Vascular morphology was monitored weekly using epifluorescence microscopy (A), Scale bar, 500 μm. Confocal immunofluorescence imaging was performed at selected timepoints to evaluate tight junction and endothelial markers (e.g., CD31, Claudin-5) (B), Scale bar 200 μm. Western blot analysis was used to assess protein expression levels of Claudin-5 (C) and CD31 at week 10 (D). These data represent three independent experiments with two to four biological replicates per condition. Values are presented as mean ± SEM. *P < 0.05, **P < 0.01 by one-way ANOVA with Tukey’s post hoc test.

The comparison of protein expression levels between density conditions was evaluated using Western blot analysis of proteins. Figure 1C depicts CD31 protein expression (CD31 expression in target groups was normalized to 0.5 million hBMEC per scaffold condition). For the 1 million hBMEC per scaffold seeding condition, CD31 levels did not differ at 2 or 4 weeks but decreased significantly by 10 weeks (P<0.05). For the 2 million cells group, CD31 levels were non-significantly reduced at 2 weeks compared to the 0.5 million group, and significantly increased from 2 weeks to 4 weeks. In the 1 million hBMEC per scaffold condition, claudin-5 levels were significantly increased at 10 weeks compared to both earlier timepoints as well as the 0.5 million hBMEC per scaffold condition average (P<0.0001) (Figure 1D). In the 2 million hBMEC per scaffold condition, claudin-5 levels were significantly increased at 10 weeks compared to the 0.5 million condition average (P<0.05).

Taken together, these data indicate that endothelial cells can autonomously form short-lived vessel-like networks; however, additional support, such as extracellular matrix or neurovascular cell types, is likely required for long-term maintenance. Given its performance advantage over low-density cultures and resource efficiency compared to high-density groups, the 1 million cell condition was selected as the negative control for downstream co-culture optimization studies.

### Extracellular matrix accelerates early vascular assembly but fails to sustain long-term endothelial stability

Following the investigation of hBMEC seeding density, we next hypothesized that the extracellular matrix (ECM) plays a critical role in promoting the formation, organization, and early maintenance of brain-like vascular networks. To test this, we evaluated four ECM proteins frequently associated with cerebrovascular development and *in vitro* endothelial culture: collagen type IV (the predominant basement membrane collagen in the brain), fibronectin and laminin (two key adhesion and signaling molecules found in the brain extracellular niche), and Matrigel (a complex ECM derived from murine sarcoma, rich in laminin, collagen IV, and growth factors). As a negative control, we included scaffolds seeded with one million hBMECs and containing collagen type I without any additional ECM supplementation.

GFP+ hBMEC monocultures with extracellular matrix-supplemented (Collagen IV, fibronectin, laminin, or Matrigel) hydrogels were monitored weekly using epifluorescence microscopy to track the formation and persistence of the vascular network (Figure 2, Supplementary Fig. 2). As seen in Figure 2A, both the unsupplemented hydrogel and the Matrigel-supplemented condition exhibit vessel-like networks at 2 and 4 weeks. Confocal images, shown in Figure 2B, demonstrate the presence of elongated CD31+ and GFP+ tube-like structures at 2 weeks after seeding.

**Fig. 2:**
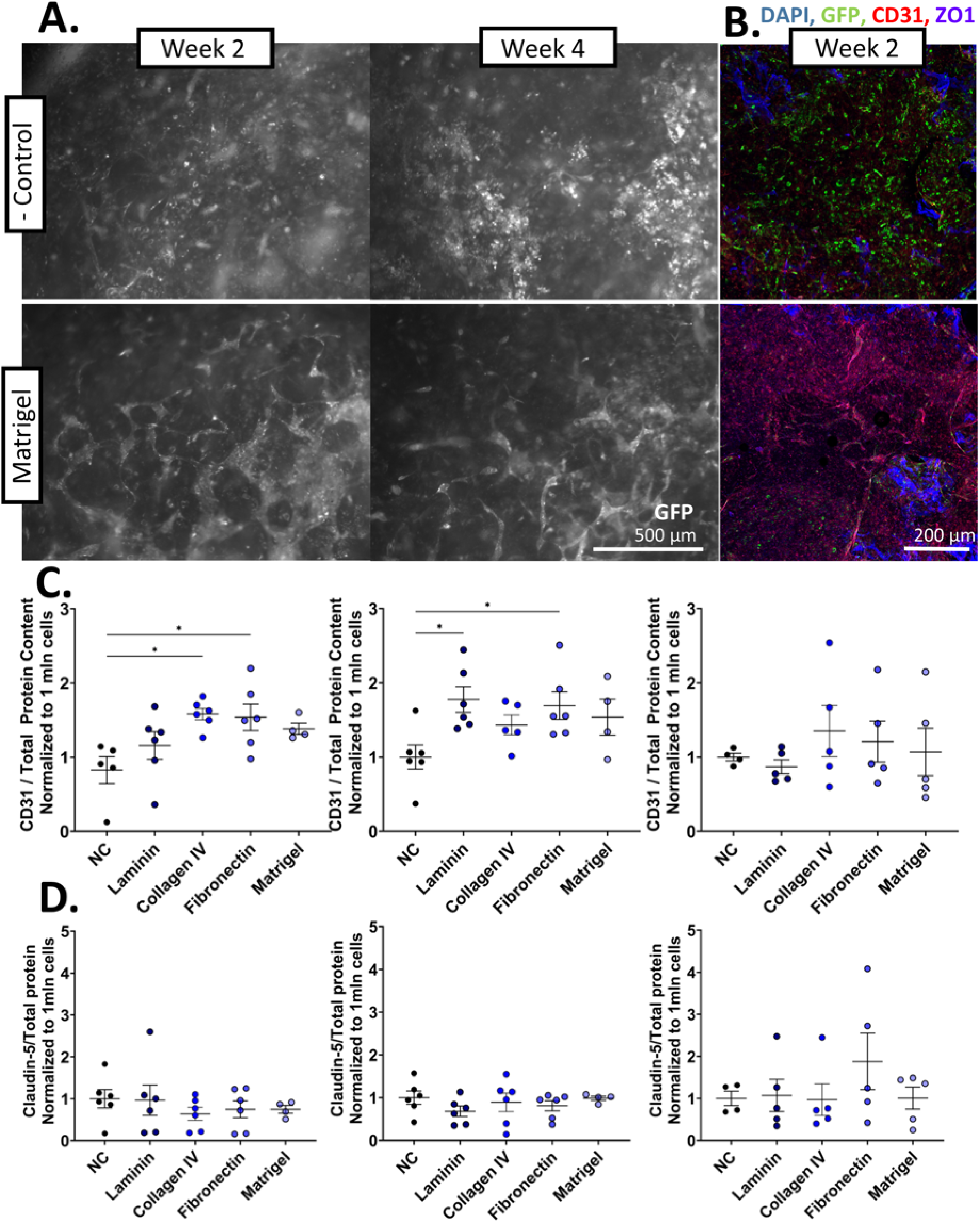
Extracellular matrix composition supports acute vascular-like network formation but does not prevent long-term structural decline in hBMEC monocultures. GFP-labeled human brain microvascular endothelial cells (hBMECs) were seeded at 1 million cells per scaffold and cultured with different extracellular matrix (ECM) supplements: collagen IV, fibronectin, laminin, or Matrigel incorporated into collagen I. Cultures without additional ECM served as negative controls. (A) Epifluorescent images of GFP-labeled hBMECs seeded with Matrigel, including or excluding ECM supplementation, at 2 and 4 weeks after seeding. Scale bar, 500 μm. (B) Confocal immunofluorescence images of scaffolds fixed 2 weeks after seeding. DAPI (405), hBMEC-produced GFP (488), CD31 (594), and ZO1 (647). Scale bar, 200 μm. (C) CD31 comparison of hBMEC scaffold monocultures without ECM supplementation to scaffolds supplemented with laminin, collagen IV, fibronectin, and Matrigel, normalized to the unsupplemented condition. (D) Claudin-5 comparison of hBMEC scaffold monocultures without ECM supplementation to scaffolds supplemented with laminin, collagen IV, fibronectin, and Matrigel, normalized to the unsupplemented condition. These data represent three independent experiments with two to four biological replicates per condition. Values are presented as mean ± SEM. *P < 0.05, **P < 0.01 by one-way ANOVA with Tukey’s post hoc test.

To compare protein expression levels between ECM conditions, western blot protein analysis was performed. Seen in Figure 2C, CD31 levels were quantified and normalized to the unsupplemented hydrogel condition. At 2 weeks, CD31 levels were increased in all ECM conditions and significantly increased in both collagen IV and fibronectin (P<0.05). At 4 weeks, CD31 levels were increased in all ECM conditions and significantly increased in both laminin and fibronectin (P<0.05). At 10 weeks, CD31 levels were not significantly increased in any condition. Claudin-5 levels were not significantly increased or decreased in any ECM condition at any time point (Fig. 2D).

Based on its consistent performance and composition, encompassing collagen IV, laminin, and fibronectin, Matrigel was selected as the ECM supplement for all subsequent co-culture experiments.

### Glial co-culture sustains the survival and complexity of the vascular network in engineered neurovascular tissues

After evaluating ECM effects, we next examined the co-culture of brain microvascular endothelial cells (hBMECs) with astrocytes and pericytes, key regulators of vascular maturation and homeostasis. We hypothesized that hBMEC monocultures are insufficient for the formation of durable vessel-like structures and that glial support is required for long-term stability. To test this, we established co-cultures of hBMECs with astrocytes (EA), pericytes (EP), or both (EPA), with and without Matrigel supplementation (EA[m], EP[m], EPA[m]) and maintained them for up to 10 weeks. Endothelial media were supplemented with an astrocyte growth factor to support glial viability. As ECM experiments did not identify a clearly superior condition, both Matrigel-supplemented and non-supplemented groups were retained to assess which condition best supported vascular structure and longevity. Matrigel, containing collagen IV, laminin, and fibronectin, served as a practical, comprehensive ECM choice for subsequent experiments.

Immunofluorescence revealed robust endothelial survival and sustained vessel-like networks in co-cultures, particularly in EPA and EPA[m], where organized CD31+ and Claudin-5+ structures persisted through week 10 (Fig. 3A). Western blot showed a modest but significant reduction in CD31 at week 2 in EA, EPA, and EPA[m] (P < 0.05), suggesting early network remodeling with glial support (Fig. 3B). Claudin-5 levels (Fig. 3C) remained stable across most conditions, while ZO-1 (Fig. 3D) expression was consistently upregulated in all glial co-cultures, indicating enhanced junctional maturation.

**Fig. 3:**
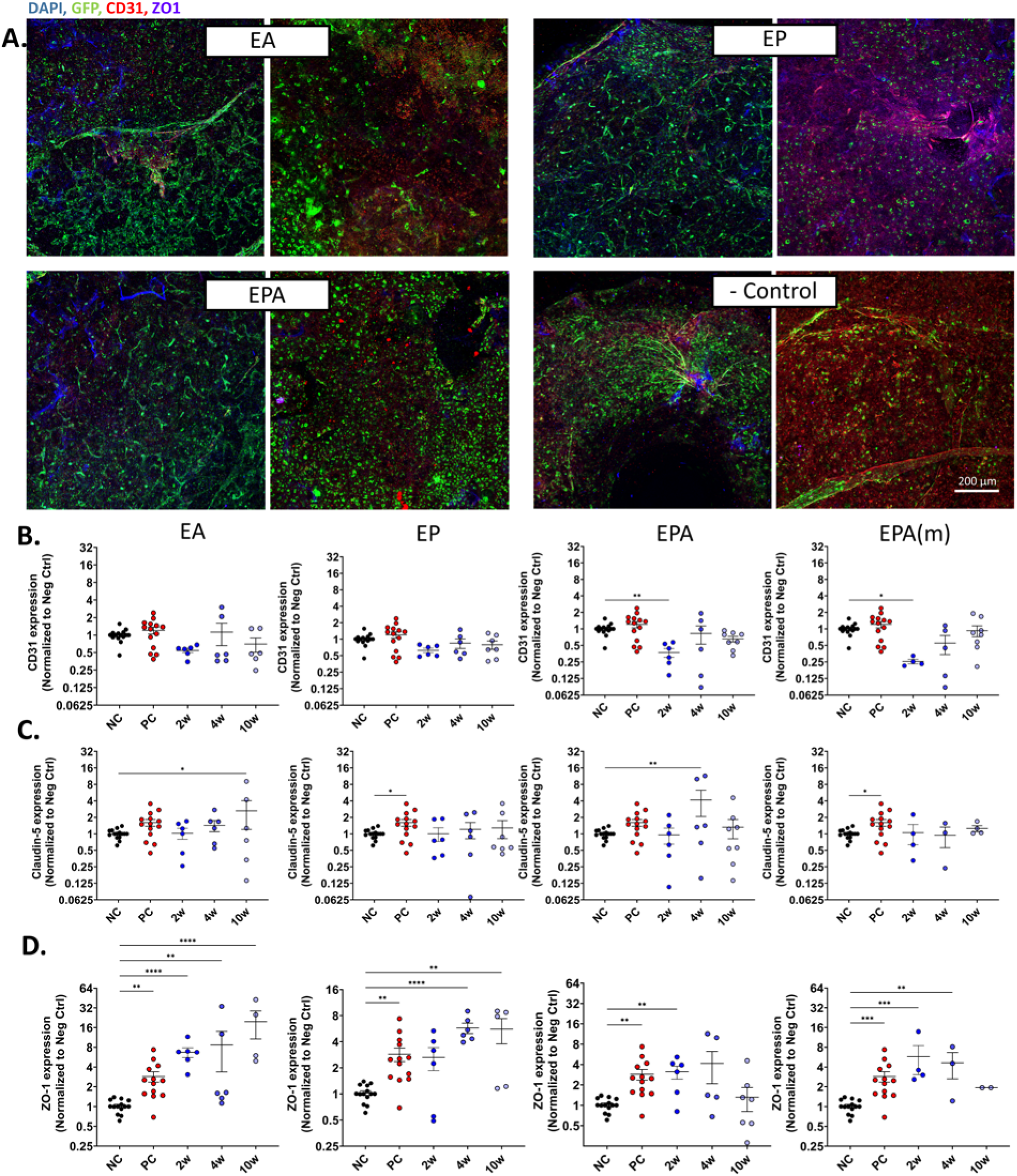
Optimization of neurovascular co-culture combinations reveals that tri-culture of endothelial cells, pericytes, and astrocytes (EPA) supports the most stable and mature vascular networks. Confocal microscopy of vascular unit co-culture. E – human brain microvascular endothelial cells, P – human brain microvascular pericytes, and A – human astrocytes. (A) Confocal immunofluorescence images of scaffolds fixed 2 weeks (left) and 10 weeks (right) after seeding for each condition. DAPI (405), hBMEC-produced GFP (488), CD31 (594), and ZO1 (647). Scale bars, 200 μm. (C-D) Western blot protein analysis of vascular unit co-cultures. E – hBMECs, P – brain microvascular pericytes, A – astrocytes, NC – negative control, hBMEC monoculture pooled across timepoints, PC – positive control, hBMEC monoculture with Matrigel pooled across timepoints. (B) CD31 comparison of NC scaffold monocultures to EA, EP, and EPA co-culture conditions, normalized to the pooled NC. (C) Claudin-5 comparison of NC scaffold monocultures to EA, EP, and EPA co-culture conditions, normalized to the pooled NC. (D) ZO1 comparison of NC scaffold monocultures to EA, EP, and EPA co-culture conditions, normalized to the pooled NC. Values are presented as mean ± SEM. *P < 0.05, **P < 0.01 by one-way ANOVA with Tukey’s post hoc test.

Morphometric analysis of the vascular network confirmed these trends: EPA[m] exhibited the highest junction density and greatest average vessel segment length by week 10 (Supplementary Figure 3A-C). Together, these findings show that astrocytes and pericytes synergistically promote vascular stability and complexity, with Matrigel further improving network organization. This approach provides a foundation for engineering durable, vascularized neural tissues for disease modeling and therapeutic development.

### Incorporation of neuronal and microglial components enhances vascular integration and network longevity in 3D neurovascular constructs

Having established that astrocytes and pericytes improve vascular stability, we next sought to recapitulate the full complexity of the human neurovascular unit by incorporating neurons and microglia into the scaffold. To this end, we extended the previously optimized triculture (EPA) model by integrating human-induced neural stem cell–derived neurons and immortalized human microglia, generating five-cell-type constructs (“PENTA” cultures). NAM triculture-seeded silk scaffolds were embedded in a 3 mg/mL collagen I hydrogel supplemented with one million GFP-labeled human brain microvascular endothelial cells (hBMECs) and 0.3 million pericytes per scaffold. Constructs were transferred to 48-well plates the following day and maintained in NAM-compatible media enriched with B27, Glutamax, Astrocyte Growth Factors, and Anti-Anti. No additional media optimization was required to support all five cell types in long-term co-culture.

As shown in Figure 4A-B, PENTA constructs developed dense, spatially organized neurovascular networks by week 5, with endothelial structures closely interlaced with neuronal and glial elements. Immunofluorescence imaging revealed colocalization of endothelial markers (CD31, ZO-1), neuronal cytoskeletal elements (TUJ1), and GFP-expressing vasculature within shared tissue domains. Western blot analysis demonstrated detectable levels of junctional proteins, including ZO-1 and CD31, confirming the persistence of endothelial identity within the multicellular context (Fig. 4B). Importantly, PENTA cultures remained viable and structurally stable for at least 16 months *in vitro*, offering a uniquely durable and human-relevant platform for studying chronic neurovascular remodeling.

**Fig. 4:**
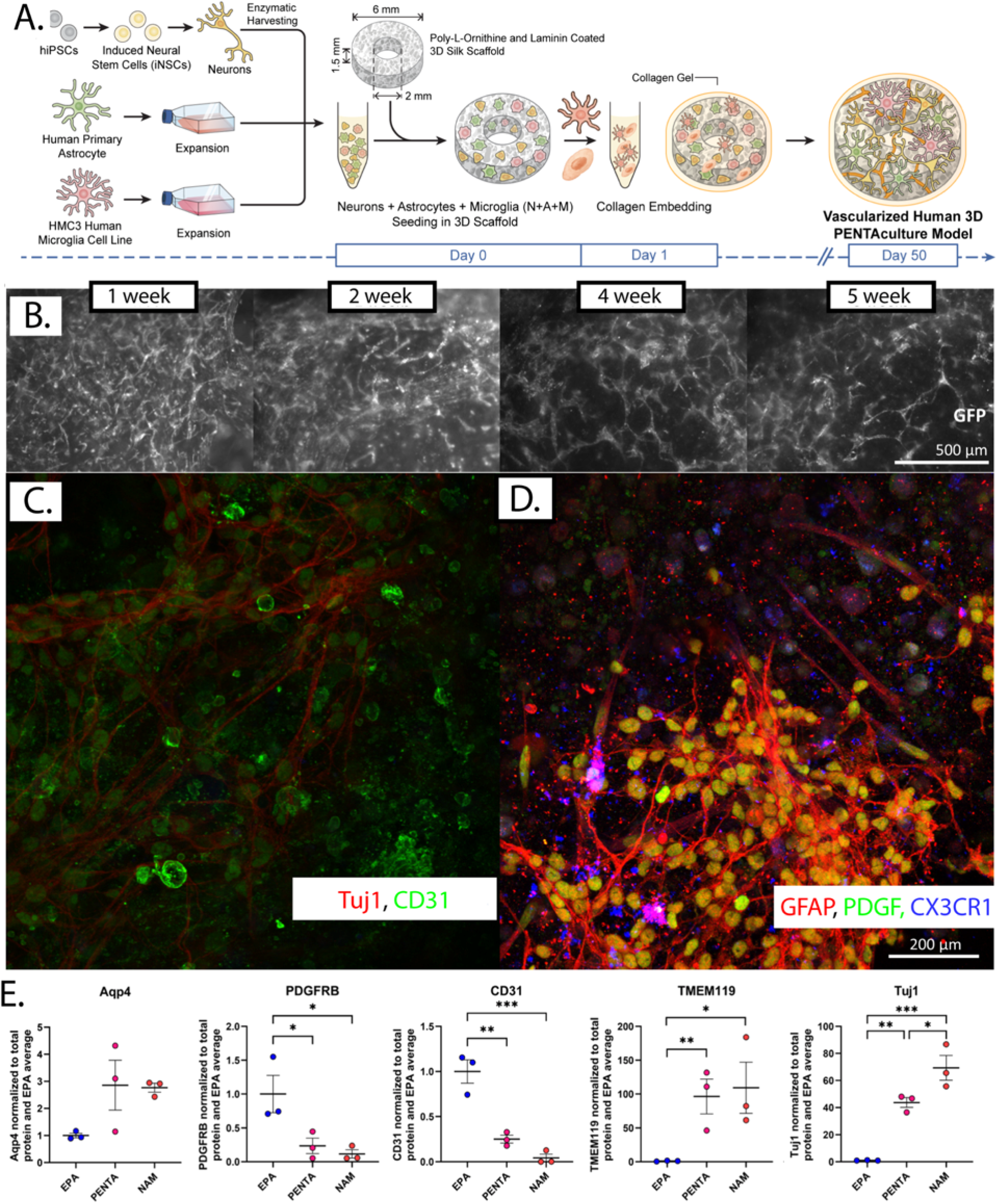
Vascularized 3D in vitro brain-like tissue (penta-cultures) model fabrication. A) Schematic Representation of PENTA-cultures fabrication; B) Representative bright field images of vascular growth over 5 weeks; C-D) Representative IF images of vascular (GFP, CD31, Z01) and neuronal networks (Tuj1), 5-weeks post-seeding of penta-cultures. E) WB results demonstrated the increase in tight junction protein (ZO-1) in penta-cultures compared to Endothelial (E)-Pericytes (P)-Astrocytes (A) triculture without neuronal population.

This system provides a structurally complex and long-lived model of the human NVU, supporting the interrogation of neurovascular interactions and injury responses with spatiotemporal resolution unmatched by simpler or shorter-lived *in vitro* approaches.

### Molecular maturation of the brain vascular unit

To evaluate the impact of neurovascular complexity on angiogenic signaling, secreted angiokines were profiled across four culture conditions of increasing cellular composition: hBMEC monocultures, hBMEC monocultures supplemented with Matrigel, EPA tricultures, and PENTA cultures, consisting of hBMECs, astrocytes, pericytes, microglia, and induced-neural stem cell (iNSC) derived neurons. As shown in Figure 5, hBMEC monocultures exhibited low or undetectable levels of pro-angiogenic factors, including VEGF-A, TGF-β1, PDGF-BB, and angiopoietins. The Matrigel condition demonstrated modest increases in select remodeling proteins such as TIMP-2 and IL-6 but did not elicit widespread angiokine expression. In the EPA co-culture condition, elevated levels of VEGF-A, PDGF-BB, MCP-1, and GM-CSF were detected. In the PENTA culture condition, broader upregulation of MMP-1, IL-10, VEGF-D, and TGF-β1 was observed, along with increased expression of angiopoietin-2 and chemokines including MCP-3, GRO, and I-TAC, suggesting synergistic neuroimmune regulation of endothelial signaling. Individual factor plots are shown in Supplementary Figures 4 and 5.

**Fig. 5:**
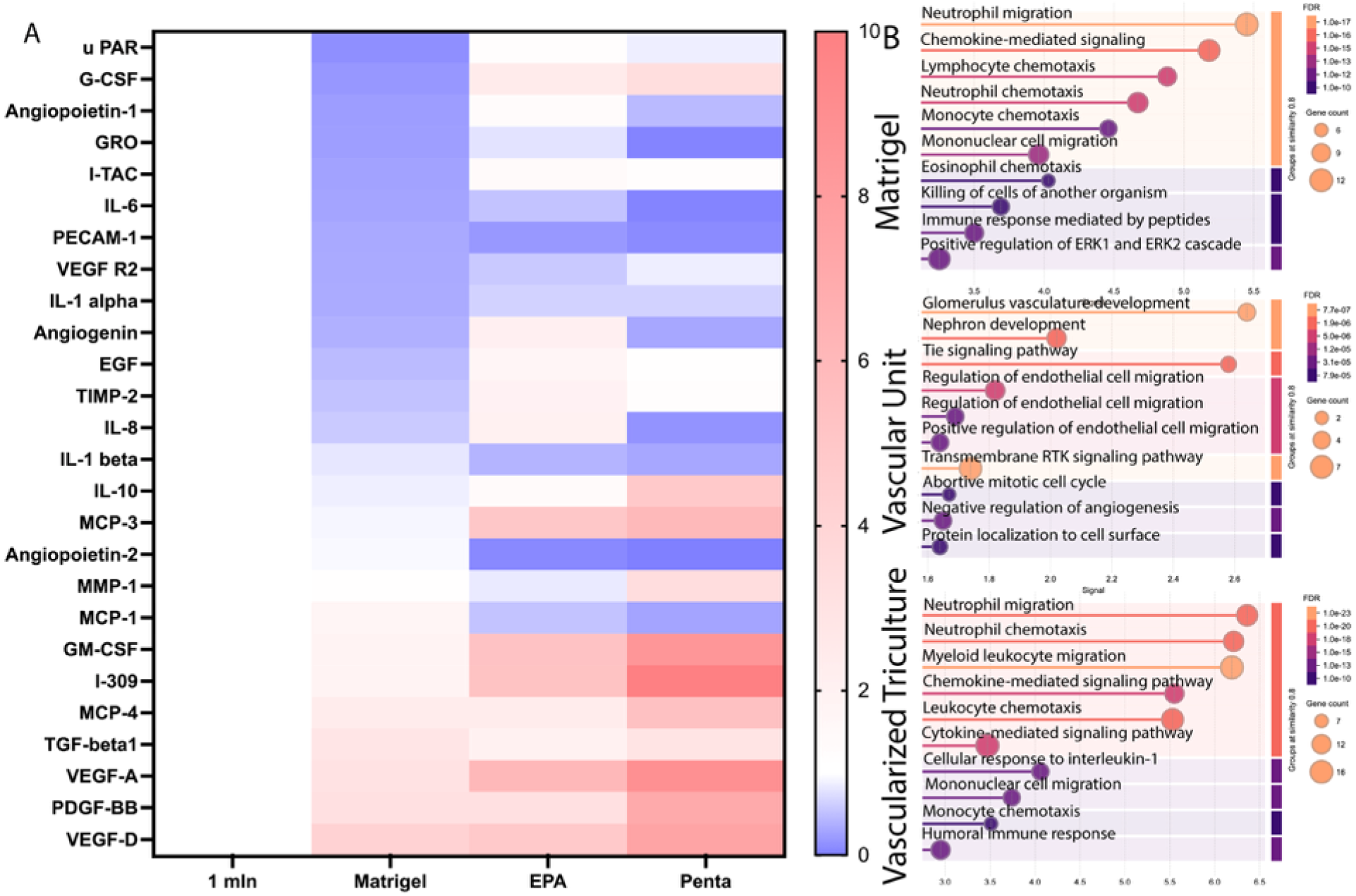
Increasing neurovascular complexity enhances angiogenic and signaling pathway activation in engineered brain tissue constructs. (A) Heatmap analysis of angiogenesis assay factors in the hBMEC monoculture (control), Matrigel, EPA, and PENTA co-cultures. Angiogenic factors measured using Raybiotech Angiogenesis Array C1000. Conditions include hBMEC monocultures (1 million hBMECs per scaffold, referred to as 1 mln), BMECs embedded in Matrigel, co-culture with astrocytes and pericytes (EPA), and a vascularized PENTA culture including neurons and microglia (PENTA). (B) Gene ontology enrichment analysis was performed to identify biological processes associated with cytokine profiles from each experimental condition. Cytokine expression data were first normalized, and differentially expressed analytes were identified. GO enrichment analysis was conducted using curated biological process annotations to assess upregulated and downregulated pathways related to vascular development, immune signaling, and cellular communication. N=4 membranes per condition with technical duplicate markers on each membrane and at least 2 experimental replicates pooled per membrane.

Pathway enrichment analysis confirmed a stepwise activation of angiogenic and endothelial-specific processes across the complexity gradient (Fig. 5B). While Matrigel cultures modestly engaged transmembrane receptor serine/threonine kinase and SMAD pathways, these were significantly enriched only in EPA and PENTA conditions. PENTA cultures exhibited selective enrichment in pathways associated with vasculature development, protein localization to the cell surface, and positive regulation of angiogenesis, alongside suppression of immune activation programs including leukocyte migration, cytokine response, and neutrophil chemotaxis. These findings suggest that neurons and microglia not only amplify angiogenic signaling in BMECs but also fine-tune the balance between vascular remodeling and inflammation. Collectively, the data highlight the necessity of multicellular neurovascular interactions for accurate modeling of brain-specific angiogenesis and support the use of PENTA systems to recapitulate physiologically relevant vascular remodeling programs for disease modeling and therapeutic development.

### Spatiotemporal disorganization of endothelial structure and junctional integrity in injured vascular units

To model post-traumatic neurovascular remodeling, EPA co-cultures were subjected to controlled cortical impact (CCI) injury of moderate intensity (6 m/s impact velocity, 3 mm diameter tip, 0.6 mm depth) and evaluated at five post-injury timepoints: 48 hours, 72 hours, 7 days, 2 weeks, and 5 weeks. Injured samples (6 m/s, 3 mm impact tip), referred to as the CCI group, were compared to sham EPA co-cultures, which received an equal duration of air exposure during the injury protocol but did not undergo mechanical injury.

Immunofluorescence staining included GFP (endothelial cytoplasm), CD31 (endothelial adhesion), TOMM20 (mitochondrial membrane), and DAPI (nuclei). To characterize endothelial structural responses following traumatic mechanical injury, EPA co-cultures were imaged with confocal microscopy at 48 hours, 72 hours, 7 days, and 5 weeks post-injury, as shown in Figure 6A. At early timepoints (48 hours and 72 hours), GFP+ endothelial cells exhibited fragmented morphology and loss of tubular organization. TOMM20 staining showed fragmented mitochondrial structures, and overall intensity appeared diminished relative to sham. By 7 days post-injury, endothelial cells began forming clustered aggregates. However, vascular alignment remained disorganized based on the GFP signal. By 1 week post-injury, partial reappearance of GFP+ tubular structures and CD31 signal was observed, with similar trends continuing out to 5 weeks. Shown in Figure 6B, quantification of vascular metrics did not demonstrate significantly altered junction density, average segment length, or lacunarity at any post-injury timepoints compared to sham. However, both lacunarity and junction density showed increased variability at 48 hours post-injury.

**Fig. 6:**
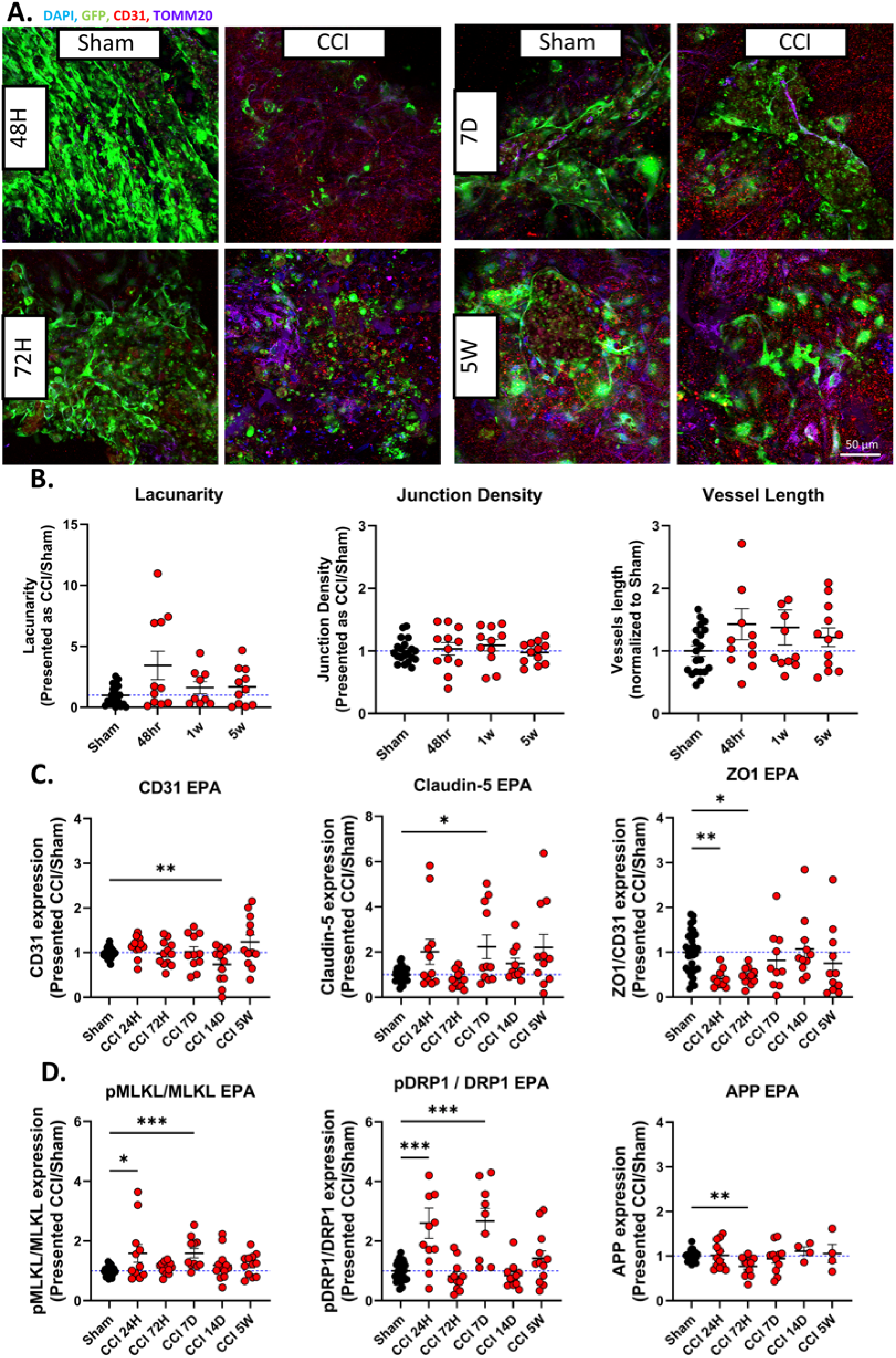
Time-resolved analysis of vascular disintegration and remodeling following moderate injury in engineered EPA neurovascular constructs. Confocal microscopy of EPA moderate injury CCI and sham samples (6 m/s impact, 3mm impact tip). A) Confocal immunofluorescence images EPA co-culture of scaffolds fixed 48 h, 72 h, 7 days, and 5 weeks after being subject to moderate injury. DAPI (405), hBMEC-produced GFP (488), CD31 (594), and TOMM20 (647). Scale bar, 50 μm. B) AngioTool quantification of vascular-like network lacunarity, junction density, and vessel length normalized to sham. (C-D) Western blot analysis of EPA moderate injury and EPA sham samples (6 m/s impact, 3mm impact tip). C) Comparison of vascular markers including CD31, Claudin-5, and ZO1. D) Comparison of mitochondrial markers, including pDRP1/DRP1 ratio and neurodegeneration-associated protein APP. Values are presented as mean ± SEM. *P < 0.05, **P < 0.01 by one-way ANOVA with Tukey’s post hoc test.

To evaluate vascular, mitochondrial, inflammatory, and neurodegenerative responses following injury, we performed western blot analysis of EPA cultures at multiple post-injury timepoints (Fig. 6C-D, Supplementary Figure 6). CD31 expression remained largely stable, with a significant decrease observed at two weeks post-injury, whereas Claudin-5 showed a transient increase at 7 days. ZO-1 expression was significantly reduced at 24 and 72 hours, consistent with early junctional disruption. Mitochondrial dynamics were altered post-injury, as evidenced by a significant increase in the pDRP1/DRP1 ratio at 24 hours and again at 7 days, indicating enhanced mitochondrial fission. TOMM20 expression remained unchanged across timepoints, suggesting preserved mitochondrial mass despite fission activity.

Next, inflammatory signaling was modest. IL-1β levels did not change significantly, whereas the phosphorylated MLKL/MLKL ratio, a marker of necroptosis, increased at both 24 hours and 7 days, indicating injury-induced necroptotic signaling. While neurodegenerative markers revealed limited change in TDP-43 expression, APP levels were more variable, elevated at 24 hours and significantly reduced by 48 hours, suggesting acute regulation following injury.

Together, these findings demonstrate that moderate injury induces sustained endothelial disorganization and junctional breakdown in EPA cultures, mimicking key features of neurovascular pathology observed *in vivo* following traumatic brain injury ^28^. The delayed or absent reconstitution of ZO-1 suggests long-term deficits in BBB-like function, reinforcing the relevance of multicellular human *in vitro* models for studying cerebrovascular dysfunction after TBI.

### Persistent pro-inflammatory and chemokine-driven signaling dominates long-term vascular response in injured EPA neurovascular cultures

To assess how limited neurovascular complexity affects angiogenic and inflammatory signaling after injury, we profiled secreted factors in EPA cultures using a RayBiotech C1000 cytokine array (Fig. 7, Individual factor plots shown in Supplementary Figures 7 and 8). At 24 hours post-injury, pathway enrichment analysis revealed downregulation of receptor tyrosine kinase signaling cascades, including MAPK and JAK-STAT pathways, alongside reduced levels of angiogenic and pro-survival mediators such as VEGF family members, PLGF, bFGF, and TGF-β1. These changes corresponded with diminished enrichment for canonical angiogenesis pathways.

**Fig. 7:**
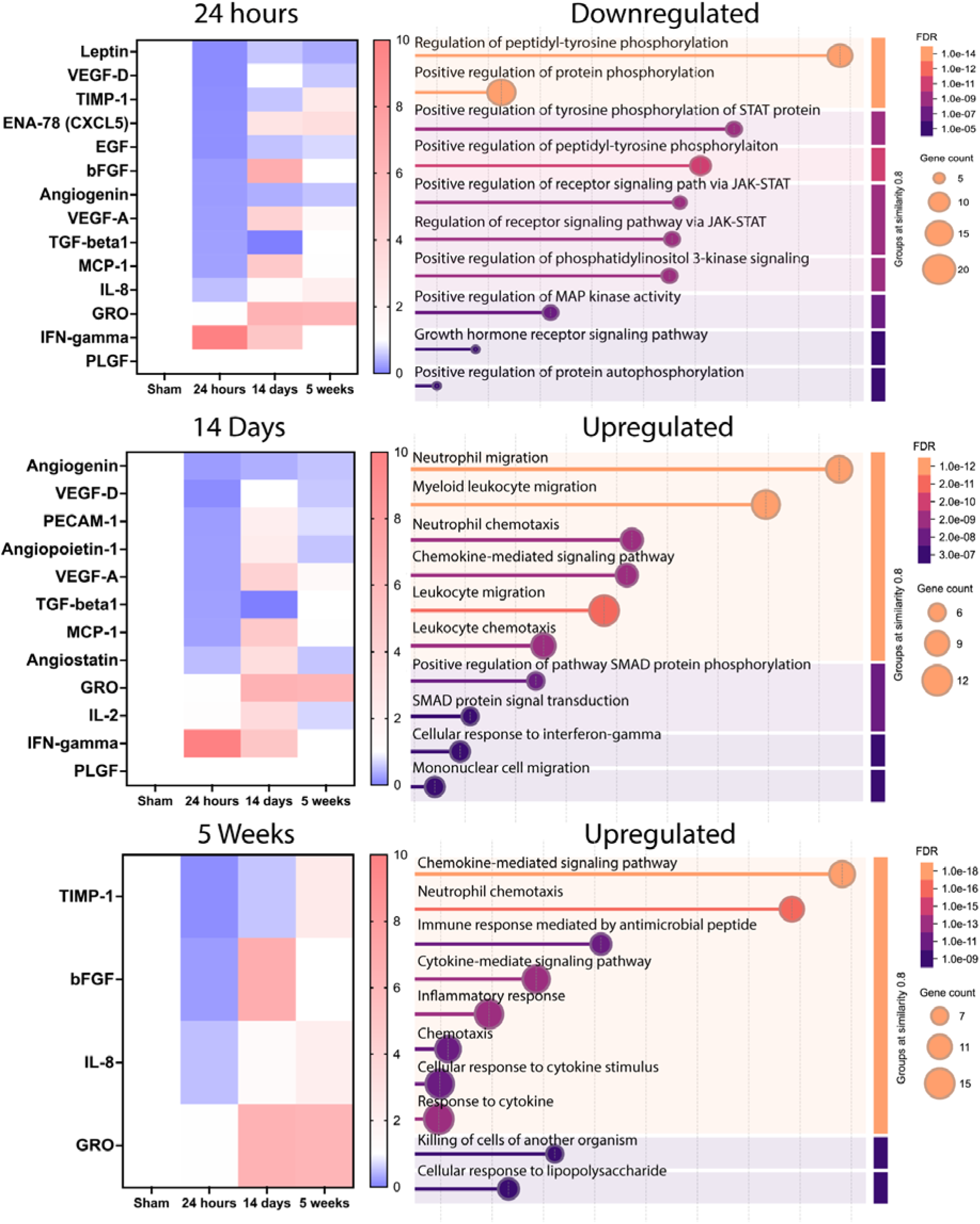
Temporal reprogramming of angiogenic and inflammatory signaling in injured EPA cultures following traumatic brain injury. Gene ontology enrichment analysis of EPA moderate injury and EPA sham samples (6 m/s impact, 3mm impact tip). Factors measured using a Raybiotech Angiogenesis Array C1000. Samples collected at 24 h and 2 weeks post-injury were normalized to sham signals. N=3 sham N=4 CCI membranes with technical duplicate markers on each membrane and 3 experimental replicates pooled per membrane. Color-coded false discovery rate (FDR) depicting gradient from purple (highest likelihood of false discovery, max 3.0 e-7) to yellow (lowest likelihood of false discovery, min 1.0 e-12).

By 14 days post-injury, the secretome shifted toward a pro-inflammatory profile, with increased secretion of MCP-1, GRO, IFN-γ, and angiogenin. Pathway analysis indicated upregulation of immune-related programs, including neutrophil and myeloid cell migration, leukocyte chemotaxis, and SMAD signaling. Despite slight increases in VEGF-A and TGF-β1, angiogenic signaling remained limited, and anti-angiogenic factors such as angiostatin and IL-2 persisted.

At last, by 5 weeks, inflammatory cytokines including GRO, IL-8, and IL-6 remained elevated, while angiogenic mediators such as bFGF and VEGF showed no reactivation. Enrichment of chemokine signaling, antimicrobial responses, and LPS-like cytokine activation suggested persistent inflammatory activity without vascular regeneration.

These findings indicate that EPA cultures exhibit sustained inflammatory remodeling without re-engagement of angiogenic programs, likely reflecting the absence of neuroimmune regulatory inputs required for resolution and repair.

### Neuroimmune interactions promote partial vascular repair but fail to resolve endothelial and mitochondrial dysfunction following brain injury

To assess how neurovascular complexity influences cerebrovascular injury responses, we subjected human PENTA cultures, comprising endothelial cells, astrocytes, pericytes, neurons, and microglia, to moderate mechanical trauma (6 m/s impact velocity, 3 mm tip, 0.6 mm depth) and analyzed structural and molecular changes at 24 hours and 14 days post-injury. Immunofluorescence revealed acute endothelial disorganization, characterized by fragmented GFP structures, reduced CD31 and ZO-1 signal intensity, and disrupted Claudin-5 expression (Fig. 8A-B, Supplementary Figure 9). By 14 days, partial recovery of endothelial architecture was observed, including reformation of GFP tubular networks and re-expression of CD31 and Claudin-5. However, ZO-1 localization remained sparse, and vessel metrics did not return to baseline, indicating persistent barrier dysfunction. APP and TDP-43 levels remained elevated, and pericyte coverage and segment length remained reduced, consistent with incomplete neurovascular repair (Fig. 8A-B, Supplementary Figure 9).

**Fig. 8:**
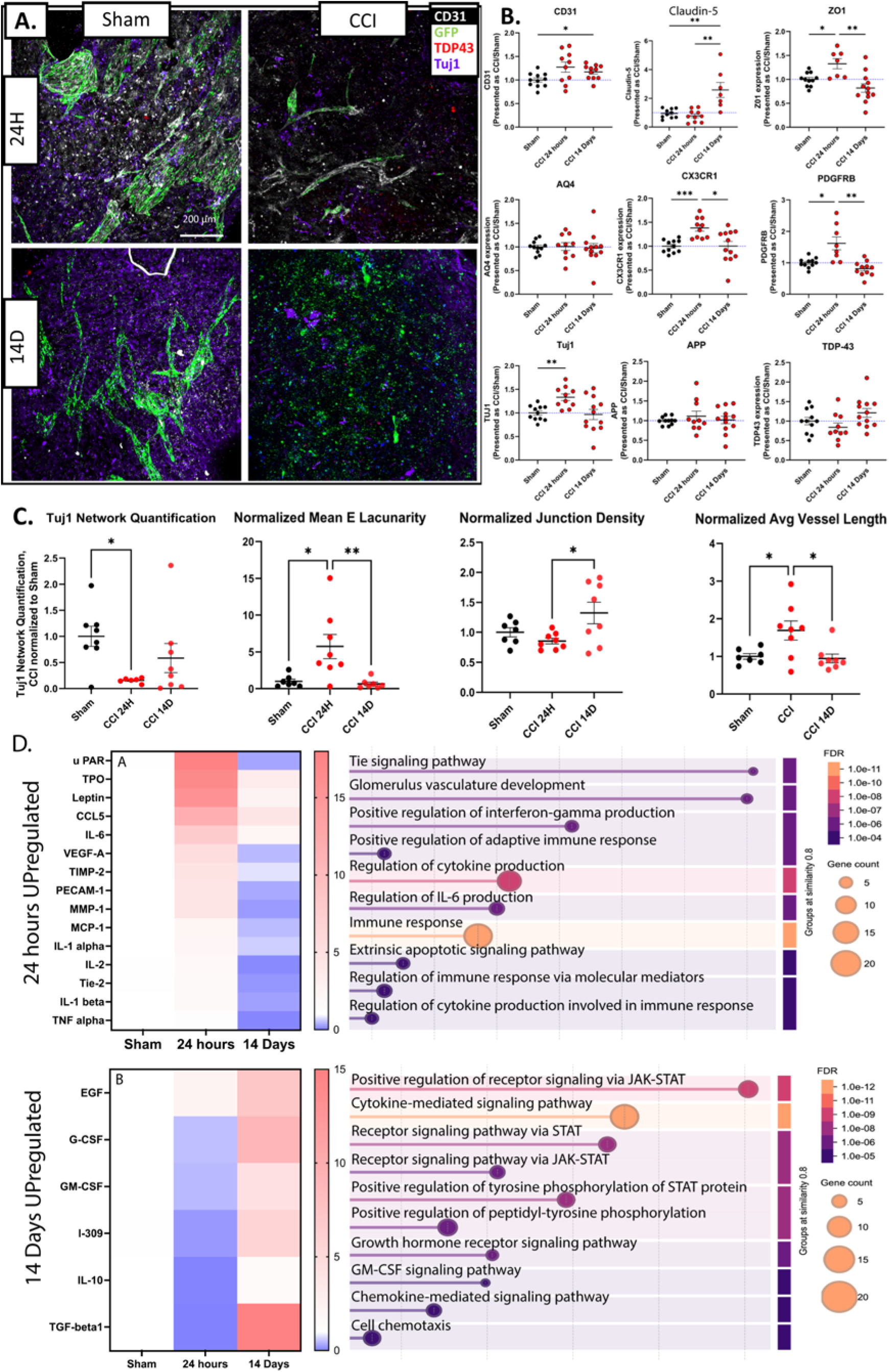
Neuronal and microglial inclusion in PENTA cultures preserves vascular integrity and shapes time-resolved inflammatory and angiogenic responses after traumatic brain injury. Confocal Microscopy and Western Blot of PENTA moderate injury and sham samples (6 m/s impact, 3mm impact tip). A) Confocal immunofluorescence images of scaffolds fixed 24 h and 14 days after being subject to moderate injury. CD31 (405), hBMEC-produced GFP (488), TDP43 (594), and Tuj1 (647). Scale bar, 200 μm. (B) Western blot analysis of PENTA moderate injury and sham samples (6 m/s impact, 3mm impact tip): vascular markers (CD31, Claudin-5, and ZO1); cell type markers (PDGFRß, CX3CR1, and AQ4); mitochondrial markers (pDRP1/DRP1 ratio and TOMM20); neuron-associated proteins (Tuj1, TDP43, and APP). (C) Tuj1 network quantification via MATLAB, normalized to sham and AngioTool quantification of vascular-like network lacunarity, junction density, and vessel length normalized to sham. (D) Gene ontology enrichment analysis of PENTA moderate injury and sham samples (6 m/s impact, 3mm impact tip). Factors measured using a Raybiotech Angiogenesis Array C1000. Samples collected at 24 hours and 2 weeks post-injury were normalized to sham signals. N=3 sham, N=4 CCI membranes with technical duplicate markers on each membrane, and 3 experimental replicates pooled per membrane. Color-coded false discovery rate (FDR) depicting gradient from purple (highest likelihood of false discovery, max 3.0 e-5) to yellow (lowest likelihood of false discovery, min 1.0 e-12). Values are presented as mean ± SEM. *P < 0.05, **P < 0.01 by one-way ANOVA with Tukey’s post hoc test.

To evaluate protein-level changes associated with vascular integrity, inflammation, mitochondrial stress, and neurodegeneration in PENTA cultures, we performed western blot analysis at 24 hours and 14 days post-injury. Vascular markers exhibited dynamic regulation across timepoints. CD31 expression increased at both time points, with a statistically significant elevation at 14 days. Claudin-5 remained stable at 24 hours but was significantly upregulated at 14 days compared to both sham and injured cultures at 24 hours. In contrast, ZO-1 was transiently elevated at 24 hours and returned to baseline by 14 days, indicating early tight junction reorganization followed by partial stabilization.

Cell type–associated markers reflected injury-responsive shifts. Platelet-derived growth factor receptor beta (PDGFRβ), a marker of pericytes and endothelial cells, was significantly elevated at 24 hours but returned to sham levels by 14 days. CX3C chemokine receptor 1 (CX3CR1), a microglia-associated marker, showed a similar transient increase at 24 hours. Aquaporin-4 (AQP4), a water channel expressed by astrocytes, remained unchanged across timepoints. Neuronal markers displayed time-dependent modulation. β3-tubulin (Tuj1) was significantly increased at 24 hours but normalized by 14 days. Amyloid precursor protein (APP) levels were unchanged. Still, they showed increased variability, while TAR DNA-binding protein 43 (TDP-43) trended downward at 24 hours and upward at 14 days, suggesting potential early suppression and delayed accumulation of neurodegenerative markers.

At last, mitochondrial analysis revealed a significant increase in TOMM20 at 24 hours, followed by a significant drop below sham levels at 14 days, suggesting acute mitochondrial activation followed by possible loss or dysfunction. The ratio of phosphorylated dynamin-related protein 1 to total DRP1 (pDRP1/DRP1) showed elevated variability, with a non-significant trend toward increased mitochondrial fission at 14 days.

### Neuroimmune complexity drives a biphasic shift from vascular inflammation to reparative signaling following traumatic brain injury

To investigate the temporal dynamics of angiogenic and inflammatory signaling after traumatic brain injury in a multicellular human neurovascular model, we profiled the secretome of PENTA cultures, comprising endothelial cells, astrocytes, pericytes, microglia, and neurons, at 24 hours and 14 days post-injury (Fig.8, Individual factor plots shown in Supplementary Figures 10 and 11).

Analysis of secreted factors revealed a biphasic response. At 24 hours, the profile was dominated by pro-inflammatory and angiogenic mediators, including vascular endothelial growth factor A (VEGF-A), interleukin-6 (IL-6), monocyte chemoattractant protein-1 (MCP-1), matrix metalloproteinase-1 (MMP-1), C-C motif chemokine ligand 5 (CCL5), and tissue inhibitor of metalloproteinases-2 (TIMP-2), alongside increased expression of the endothelial markers Tie-2 and PECAM-1, consistent with early vascular destabilization and immune activation. Elevated levels of IL-1α, IL-1β, and tumor necrosis factor-alpha (TNF-α) further reflected an acute pro-inflammatory state. By 14 days, the secretome shifted toward a reparative phenotype. Pro-inflammatory cytokines declined, while interleukin-10 (IL-10), transforming growth factor-beta 1 (TGF-β1), granulocyte-macrophage colony-stimulating factor (GM-CSF), granulocyte colony-stimulating factor (G-CSF), and epidermal growth factor (EGF) were upregulated, indicating immune modulation and a transition toward endothelial repair and tissue remodeling.

Pathway enrichment analysis supported this progression: early responses were associated with immune activation, IL-6 signaling, and apoptosis, while later timepoints showed enrichment for JAK-STAT signaling, growth hormone and GM-CSF receptor pathways, and chemokine-mediated cell migration.

These results demonstrate that PENTA cultures recapitulate key features of post-traumatic vascular remodeling, transitioning from acute endothelial activation and cytokine-driven inflammation to a late-phase reparative program shaped by multicellular neuroimmune interactions. Individual factor plots shown in Supplementary Figure 10 and 11.

## Discussion

The development of our long-lived, human-derived PENTA-culture neurovascular model marks a significant advancement in *in vitro* modeling of the neurovascular unit (NVU). By incorporating five major cell types, brain microvascular endothelial cells, pericytes, astrocytes, microglia, and neurons, into a 3D scaffold, we have achieved a structurally and functionally complex system that captures key aspects of NVU physiology and pathology. Unlike traditional monoculture, endothelial-glia triculture, or even advanced microfluidic BBB platforms, the PENTA model enables extended co-culture beyond 10 weeks and provides high spatiotemporal resolution to study injury-induced remodeling at the cellular, molecular, and secretory levels^25,27,29,30^.

A central advance of this model is its capacity to dissect chronic neurovascular dysfunction following traumatic brain injury (TBI) with cell-type specificity. Our findings reveal that moderate mechanical injury elicits a biphasic response in PENTA cultures: an early phase of endothelial disintegration, mitochondrial stress, and glial activation, followed by a delayed and incomplete attempt at repair. These temporal patterns of protein expression and structural changes suggest coordinated but ultimately insufficient restoration of neurovascular architecture and barrier function.

Importantly, our cytokine profiling revealed injury-induced activation of chemokine-mediated signaling, JAK/STAT, and Tie2-associated angiogenic pathways. These molecular signatures closely align with *in vivo* human and animal models of TBI and neurodegeneration ^31–35^. However, such signaling has not been previously captured *in vitro* within an integrated NVU context. Enrichment of these pathways in our model, alongside sustained secretion of immune-regulatory cytokines (G-CSF, IL-6, CXCL5), underscores the role of glial-endothelial and pericyte-microglia cross-talk in shaping vascular remodeling and immune modulation. These results position the PENTA platform as a uniquely human-relevant and mechanistically rich testbed for neurovascular injury and repair.

Relative to existing models, our PENTA-culture system bridges a critical gap in NVU biology by recapitulating both injury-responsive structural phenotypes (e.g., junctional fragmentation, altered lacunarity) and chronic signaling patterns observed in patient CSF or brain tissue ^35–40^. Prior triculture BBB models, though valuable, lack neurons and microglia, two essential contributors to neurovascular crosstalk ^29^, longevity, and a full three-dimensional environment ^27^. Additionally, while models using flow systems enhance tight junction expression under physiological shear stress, they are limited in culture duration and cell-type diversity ^41,42^. Our model, while static, incorporates dynamic elements such as mechanical injury, glial reactivity, and metabolic stress that shift the system into a remodeling state, offering a complementary and highly adaptable platform.

The modularity of the PENTA system also enabled direct comparison to simpler configurations used previously in published studies ^25,27,30^. While endothelial and EPA cultures exhibited prolonged inflammatory signaling and failed revascularization, inclusion of neurons and microglia in the PENTA model enhanced junctional recovery and shifted the secretome toward a pro-regenerative, immune-modulated profile. This underscores the importance of cellular complexity in achieving tissue-level repair. Furthermore, we identified previously unreported pathway activations, namely, Tie2 and JAK/STAT signaling, as key mediators of injury-induced angiogenesis and immune crosstalk in the human NVU.

While the PENTA system offers substantial advantages in modeling chronic neurovascular remodeling, several limitations have to be considered. First, the lack of continuous fluid flow and physiological shear stress likely limits the full maturation of tight junctions and functional barrier properties. Fluid-induced shear has previously been shown to affect endothelial maturation and barrier function ^43–45^. This may explain the lower baseline expression of ZO-1 and occludin compared to flow-based NVU models. Second, the absence of circulating immune cells (e.g., monocytes, lymphocytes) constrains the system’s ability to model secondary immune infiltration and chronic neuroinflammation. Finally, the current analysis focused on protein-level and morphological readouts; future work will integrate spatial transcriptomics and single-cell RNA sequencing to define cell-type-specific responses and molecular pathways driving neurovascular degeneration and repair.

Looking forward, the modularity of the PENTA platform supports systematic interrogation of specific cellular or molecular components. Integration of perfused microchannels or dynamic flow systems could enhance endothelial maturation and enable assessment of barrier function which are known to be affected by fluid-induced shear ^43–45^. Incorporation of patient-derived iPSC lines, particularly from individuals with neurovascular or neurodegenerative disorders, could provide personalized insights into disease susceptibility and therapeutic response. Moreover, targeted perturbations of signaling pathways (e.g., JAK/STAT, VEGF, or Tie2) or cell-type interactions (e.g., pericyte–microglia communication) may help elucidate drivers of vascular stabilization versus degeneration.

In conclusion, the PENTA-culture model constitutes a transformative tool for studying human neurovascular function and pathology. It captures key mechanistic events, from vascular disintegration to incomplete repair, seen in clinical neurotrauma and neurodegeneration. Its capacity to reveal previously inaccessible glial-vascular signaling and chronic remodeling pathways holds promise for therapeutic discovery, particularly for diseases involving vascular compromise, neuroinflammation, and long-term barrier disruption.

## Ethics Declaration

### Ethics

This work did not include any experiments with animals or the participation of human subjects.

## Authors contribution

VL conceived the project. DSH, SJ, and VL designed and interpreted the experiments, and DSH and VL wrote the manuscript. DSH and SJ performed scaffold seeding and sample maintenance (media changes). DSH and SJ performed all in vitro experiments, injuries, and DSH performed Raybiotech assays. VL performed confocal imaging. DSH, SJ, CCA, and MDT conducted Western Blots. VL performed ELISA. DSH, SJ, CCA, and MDT prepared silk materials.

## Acknowledgements

Cincinnati Children’s Hospital Bioimaging and Analysis Facility.

## II. Methods

### 3D Silk scaffold preparation

Scaffolds were fabricated using silk fibroin solution extracted from Bombyx mori cocoons, following previously established protocols [6]. Briefly, the silk cocoons were boiled in a sodium carbonate solution for 30 minutes to remove sericin from the fibroin. The resulting fibers were dried overnight in a chemical fume hood, then dissolved in 9.3 M lithium bromide solution. This solution was subsequently dialyzed in deionized water for three days to remove the salt. A 6% (w/v) silk fibroin solution was then cast into a round 6 cm diameter dish and mixed with sodium chloride particles ranging in size from 300 to 425 μm. After incubating for two days, the mixture was heated at 65°C for one hour to induce beta-sheet formation, resulting in a three-dimensional sponge. The sponge was removed from the dish and dialyzed at room temperature for two additional days to leach out the sodium chloride. Scaffolds were then cut using circular biopsy punches (McMaster-Carr, Princeton, NJ, USA) to a final size of 6 mm outer diameter and 2 mm inner diameter, and trimmed to a height of 1.5 mm using a razor blade. This created cylindrical scaffolds of 6 mm outer diameter, a 2 mm interior window, and 1.5 mm of height. The scaffolds were sterilized by autoclaving for 20 minutes and stored at 4°C for up to one week prior to use.

### Multicellular engineered brain-like tissue preparation

The protocol for generating the 3D multicellular engineered brain microvascular tissue model was adapted from the Liaudanskaya laboratory’s previously published 3D human iPSC-derived cortical neuron model [6]. Silk scaffolds, created as previously described, were coated with 0.1 mg/mL poly-L-lysine (PLL) at 37°C overnight, followed by a second overnight coating with 50 μg/mL laminin at 4°C. The next day, scaffolds were washed five times with phosphate-buffered saline (PBS), each wash lasting 5 minutes. Scaffolds were then conditioned with either neural basal (NB for PENTA cultures) or endothelial medium (for hBMEC monocultures or EPA cultures) until the isolated cells were ready for seeding.

Immediately prior to seeding, scaffolds were transferred to 96-well plates, and all residual liquid was removed using a vacuum aspirating pipet. A 40 μL suspension containing cells at their respective densities (50 million cells/mL of neurons and 12.5 million cells/mL of astrocytes) in neurobasal or endothelial media was then applied to each dried scaffold and incubated at 37°C for 30 minutes to facilitate cell attachment. Following this, 150 μL of either neurobasal or endothelial medium was added to each well, and the plates were placed in a tissue culture incubator (37°C, 5% CO, humidified atmosphere).

The next day, the cell-seeded scaffolds were transferred to fresh 96-well microplates, and a collagen type I hydrogel containing endothelial cells and pericytes was added to each scaffold (100 μL of 3 mg/mL collagen solution and pH adjusted to 7.0 using NaOH. 10 million cells/mL of hBMECs and 3 million cells/mL of pericytes). After a 30-minute incubation at 37°C to allow collagen crosslinking, 150 μL of either neurobasal or endothelial medium was added to each scaffold and incubated for 24 hours. On the following day, the seeded scaffolds were transferred to 48-well plates containing 1 mL of neurobasal or endothelial medium. To support glial viability, endothelial media was supplemented with an astrocyte growth supplement in EA co-cultures and EPA tricultures. Media was refreshed every four days until the constructs reached the assigned timepoint for fixing / freezing (2, 4, or 10 weeks for characterization) or until they were subjected to controlled cortical impact (CCI) 5–6 weeks post-seeding.

### Mechanical contusion injury model

Scaffolds were mechanically injured to simulate moderate TBI through the use of a controlled cortical impactor (CCI) device. Scaffolds were placed on sterile weigh boats within a biosafety cabinet and subjected to mechanical injury using a pneumatic CCI device in a manner similar to our group’s previously published model [6]. For the moderate injury paradigm, a 3 mm diameter impactor tip was employed at a velocity of 6 m/s, with an impact depth of 0.6 mm and a dwell time of 200 milliseconds. Injuries were administered 5 weeks after scaffold seeding, with each injured sample receiving 1 hit. Sham controls were exposed to identical environmental conditions, including air exposure on the weigh boat, but did not undergo mechanical impact. Sample collection timepoints were defined relative to the injury event.

### Western blot

To compare relative levels of protein expression between groups, a Western blot was utilized. For protein extraction, whole scaffolds previously stored at −80°C were lysed by the addition of 1X RIPA buffer. Samples were then subjected to sonication at 20% amplitude for 20 pulses (1 second on, 1 second off) to facilitate protein release. The resulting lysates were diluted in a 3:1 ratio with sample buffer composed of Laemmli buffer and β-mercaptoethanol (9:1). In addition to these lysates, mitochondrial and total protein samples were collected following Seahorse assay post-processing. For electrophoresis, 15 µL of mitochondrial or sonicated scaffold lysates were mixed with 5 µL of sample buffer. All samples were denatured at 95°C for 5 minutes prior to loading onto precast acrylamide gels.

Protein separation was performed by SDS-PAGE at 150 V for approximately 50 minutes, or until the tracking dye reached the bottom of the gel. Precision Plus Protein™ All Blue Standard (Bio-Rad) was used as a molecular weight reference (5 µL for general samples; 10 µL for Seahorse protein samples). Gels were imaged using the Stain-Free setting on a Bio-Rad imager. If sample number exceeded the capacity of a single gel, samples were divided across multiple gels. For consistency, gels were sectioned at identical molecular weight markers (75 kDa and 20 kDa) prior to transfer to polyvinylidene fluoride (PVDF) membranes.

Protein transfer was performed using a semi-dry transfer system at 2.5 V and 1.6 A for 4 minutes (total protein samples) or 3 minutes (mitochondrial samples). Following transfer, membranes were imaged using the stain-free preset to assess total protein content for normalization purposes. Membranes underwent multiple rounds of antibody probing, imaging, and stripping to enable detection of various target proteins. Primary antibody incubation was carried out overnight at 4°C on a rocker using typical dilutions of 1:1000 in Blocking Buffer, or 1:3000 for particularly strong signals. Membranes were washed using Tris-buffered saline with Tween 20 (TBST), beginning with two immediate rinses, followed by three 10-minute washes. Secondary antibodies were applied at a 1:10,000 dilution in Blocking Buffer for 1 hour at 4°C, followed by the same wash sequence. Detection was performed using an enhanced chemiluminescence (ECL) substrate, with membranes exposed for 5 minutes prior to imaging. To enable subsequent rounds of probing, membranes were stripped using Western blot stripping buffer at room temperature for 30 minutes, followed by the standard TBST wash protocol.

For analysis, all markers were normalized to the total protein content of their respective samples lane. Tight junction proteins are presented as (TJ protein / CD31) / total protein content. pMLKL / MLKL ratios are presented as (pMLKL / MLKL) / total protein content. pDRP1 / DRP1 ratios are presented as (pDRP1 / DRP1) / total protein content. All other markers are presented as normalized to the total protein content.

### Angiogenesis Assay

The RayBiotech C1000 Angiogenesis Assay was used for pathway-level analysis by measuring 43 specific angiogenic factors. The protocol was followed as listed in the product specifications. Briefly, pre-loaded membranes with antibodies were coated in 200-300 µg of scaffold protein lysate, with 1 mL of solution per membrane. After incubation with primary and secondary antibodies, a detection buffer was used to quantify each target factor via chemiluminescence. Factors were compared to the Gene Ontology library and assessed using STRING protein-protein interactions ^46,47^.

### Confocal Microscopy

To visualize cellular morphology and localization of proteins, confocal microscopy and immunofluorescence were used. 3D cell-seeded scaffolds were fixed in a solution containing 4% paraformaldehyde (PFA; Electron Microscopy Sciences) and 4% sucrose in phosphate-buffered saline (PBS; Thermo Fisher) for 1 hour at room temperature. Following fixation, samples were washed with PBS and permeabilized for 1 hour using a solution of 0.2% Triton X-100 and 4% goat serum (Thermo Fisher). Primary antibody incubation was carried out overnight at 4°C in the same permeabilization solution. The following day, samples were gently washed five times with PBS before being incubated with fluorophore-conjugated secondary antibodies for 1 hour at room temperature. Nuclear staining was then performed using DAPI (4′,6-diamidino-2-phenylindole) for 5 minutes. Excess antibodies and DAPI were removed by five additional PBS washes. Fluorescent image stacks of the 3D tissues were acquired using a Zeiss confocal microscope. Regions of interest were chosen blinded to the channel of quantification (avoiding GFP, CD31, and Tuj1). 50 z-stacks were created at a 10x zoom, a resolution of 1024×1024 pixels (3181.98 µm x 3181.98 µm), and with a step size of 1.5 µm. Data presented as max projection of z-stacks with combined channels. Images were uniformly altered for visual presentation in figures only, not before any analysis.

Tuj1 neuronal network density was analyzed using a custom MATLAB code previously published by the Liaudanskaya research group ^48^. Briefly, Z-stacks of Tuj-1–labeled 3D tissues were imported into MATLAB for analysis. Background fluorescence from silk and cell bodies was removed using Otsu’s thresholding and an eccentricity filter (≥0.90) to exclude circular structures, preserving elongated, tube-like neurites. Connected neurite voxels (defined by 18-point connectivity; voxel dimensions = 0.258 μm × 0.258 μm × 0.422 μm) were grouped into individual objects. For each object, the longest principal axis of its best-fit ellipsoid was used to quantify neurite length. Neural network prevalence within each stack was calculated as the percentage of voxels included in neurite length analysis relative to the total voxel count.

Vascular-like network parameters were quantified using AngioTool, an open-source software developed by the NIH for analysis of angiogenesis^49^. All images used for analysis were maximum intensity projections of the GFP channel and included vessels on a dark background, compatible with AngioTool’s segmentation pipeline. Images were processed using a multiscale Hessian-based enhancement filter to identify tube-like structures, followed by skeletonization for quantitative morphometric analysis. Metrics computed included total and average vessel length, vessel area, number of junctions, vascular density, lacunarity, and branching index (branch points per unit area). Lacunarity, a measure of structural heterogeneity, was calculated using a fast box-counting algorithm. To ensure consistency, identical scale and segmentation parameters were applied across all groups. AngioTool’s processing leveraged Java-based libraries and ImageJ plugins, allowing reproducible, high-throughput quantification across multicore systems.

### Statistical analysis

Statistical analysis was performed using GraphPad Prism 7. Two tailed t-tests were used to compare two experimental groups. One-way ANOVA analysis of variance was used to compare multiple groups within a single timepoint or one group across multiple timepoints. Tukey’s post-hoc test was used to evaluate significant differences between experimental groups and for comparison to controls. P ≤ 0.05 was considered statistically significant. All experiments were repeated three times to minimize *in vitro* variance. At the same time, due to the independent nature of each scaffold during culture, each scaffold was treated as an independent sample. Samples were normalized to the average of the negative control from their respective experimental replicate, then all experiments were combined for statistical analysis. For example, in Figure 2, corrected western blot data (CD31 normalized to total protein content on the sample lane) were divided by the average of the 0.5 million group within each experiment. All quantified data presented are the average and standard error of the mean of three independent experiments.

