## Supplementary for "A long-lived Human Neurovascular PENTA Culture Model Reveals Incomplete Vascular Repair and Glial-Mediated Signaling After Traumatic Brain Injury"

Appendix

### Supplementary Figures

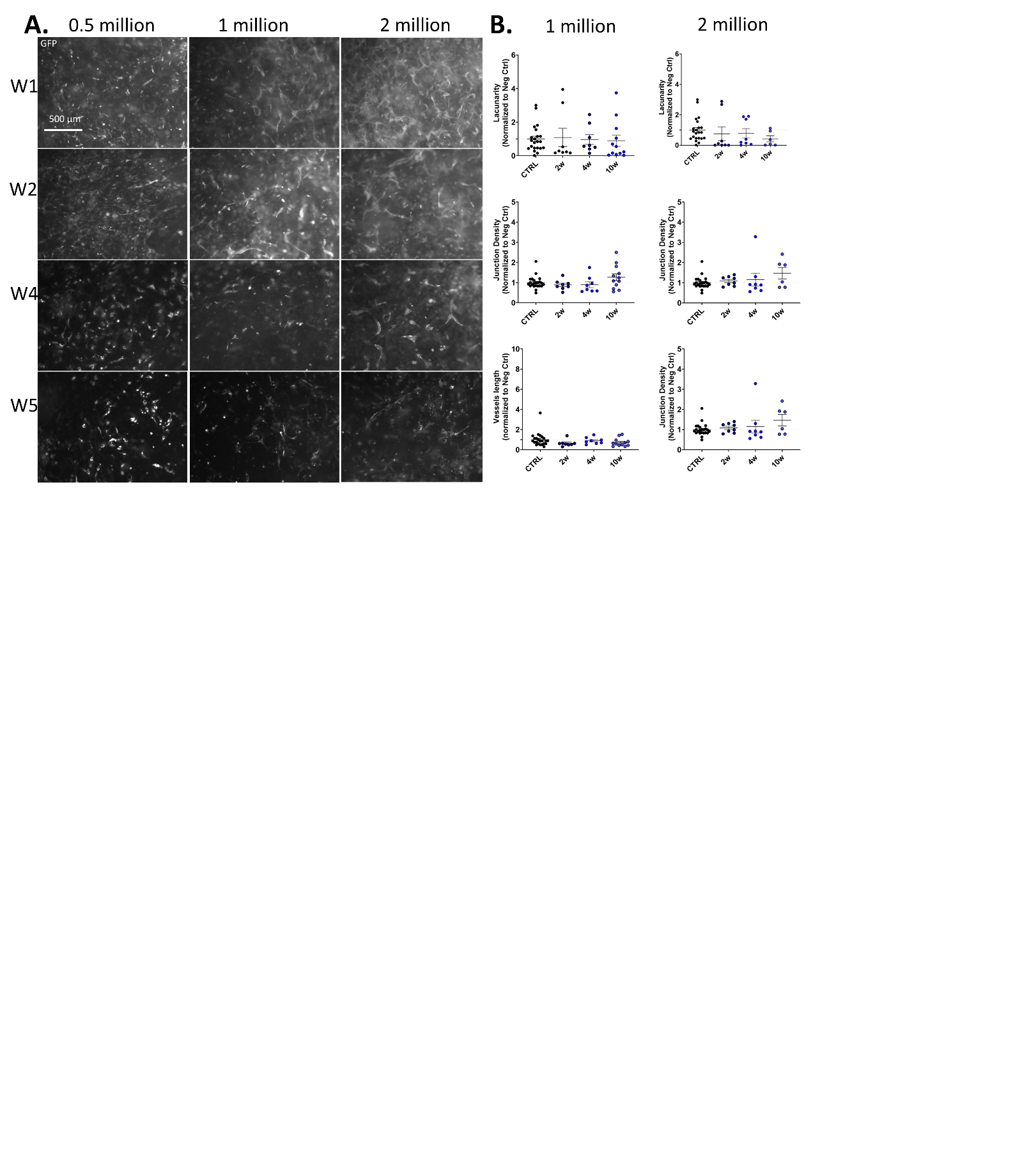

Supplementary Figure 1 – GFP-labeled human brain microvascular endothelial cells (hBMECs) were seeded at 0.5, 1, or 2 million cells per scaffold and cultured. The 0.5 million condition served as the negative control. (A) Epifluorescent images of GFP-labeled hBMECs seeded at 0.5, 1, or 2 million cells per scaffold at 2 and 4 weeks after seeding. Scale bar, 500 μm. (B) Angiotool analysis showing mean e lacunarity, junction density, and vessel length. The 1 and 2 million cell / scaffold conditions were normalized to the average of the 0.5 million condition for each metric.

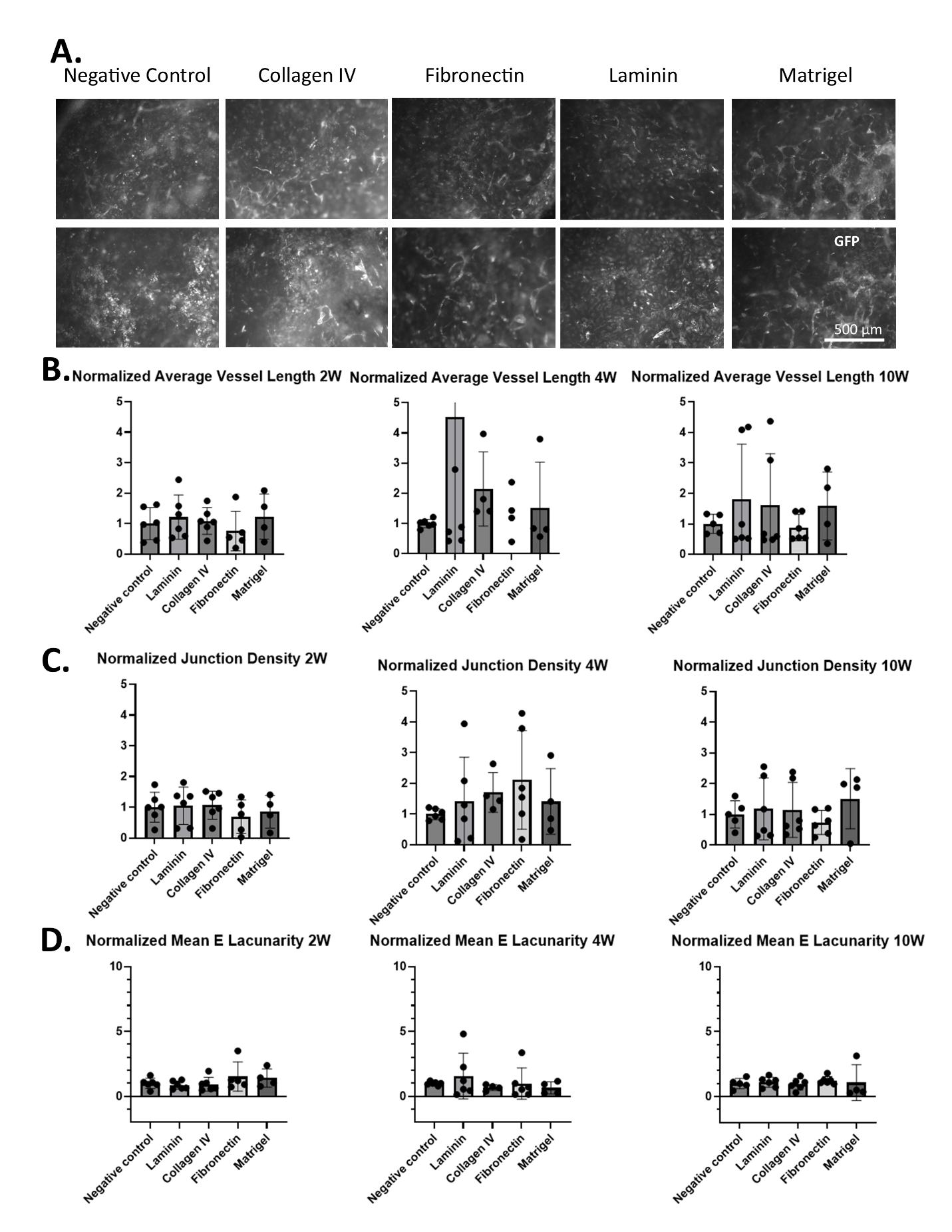

Supplementary Figure 2 – GFP-labeled human brain microvascular endothelial cells (hBMECs) were seeded at 1 million cells per scaffold and cultured with different extracellular matrix (ECM) supplements: collagen IV, fibronectin, laminin, or Matrigel incorporated into collagen I. Cultures without additional ECM served as negative controls. (A) Epifluorescent images of GFP-labeled hBMECs seeded with ECM supplementation at 2 (top) and 4 (bottom) weeks after seeding. Scale bar, 500 μm. (B) Angiotool analysis showing mean e lacunarity. (C) Angiotool analysis showing junction density. (D) Angiotool analysis showing average vessel length. The ECM-supplemented conditions were normalized to the average of the unsupplemented condition for each metric.

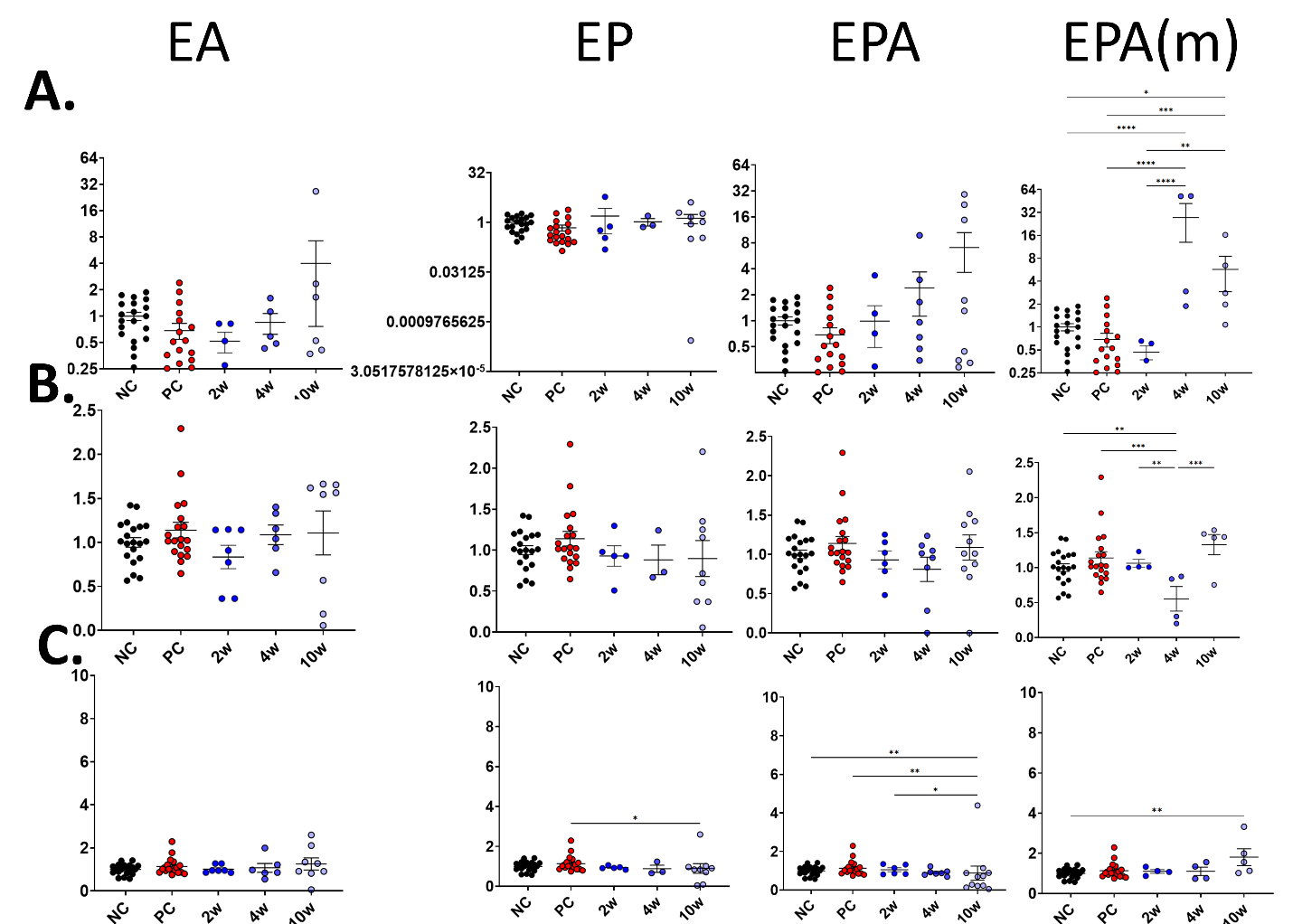

Supplementary Figure 3 – Co-culture confocal image analysis via Angiotool. E – human brain microvascular endothelial cells, P – human brain microvascular pericytes, and A – human astrocytes. (A) Angiotool analysis showing mean e lacunarity. (B) Angiotool analysis showing junction density. (C) Angiotool analysis showing average vessel length. The ECM-supplemented conditions were normalized to the average of the unsupplemented condition for each metric.

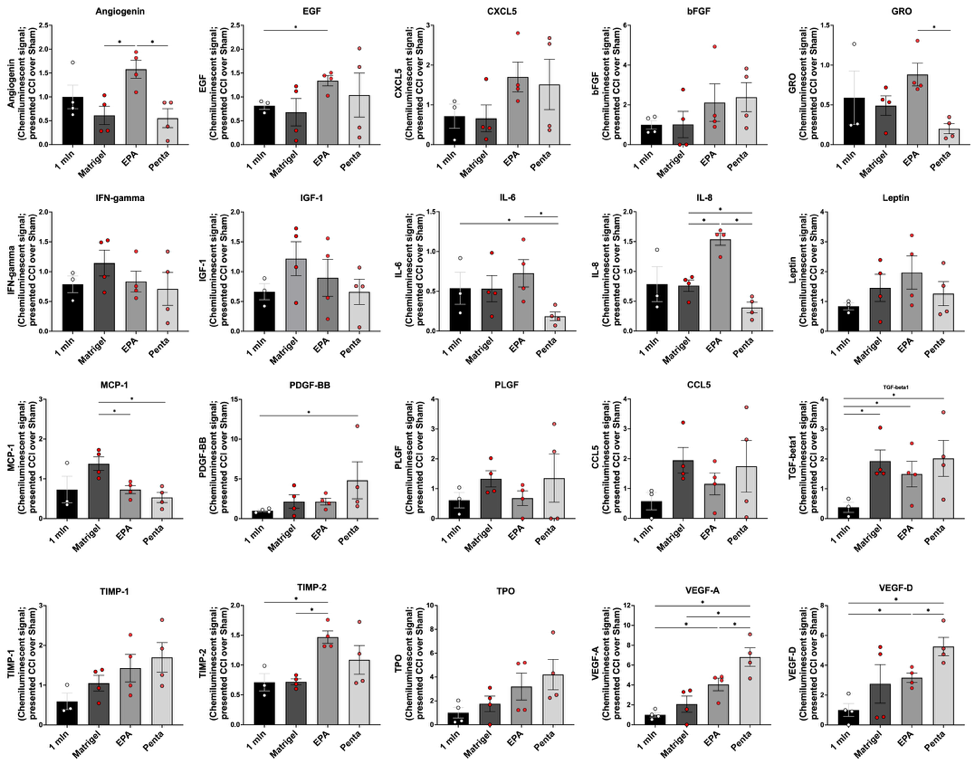
Supplementary Figure 4 - Multiplex quantification of angiogenic and inflammatory mediators across increasing vascular complexity. Bar graphs depict concentrations of 24 angiogenic and immunomodulatory proteins measured in conditioned media collected from cultures of brain microvascular endothelial cells (BMECs) alone (1 mln), Matrigel-embedded BMECs, EPA co-cultures (endothelial cells, pericytes, astrocytes), and PENTA cultures (EPA + neurons and microglia). Factors profiled include classical angiogenic mediators (VEGF-A, VEGF-D, PLGF, bFGF, PDGF-BB), extracellular matrix regulators (TIMP-1, TIMP-2), cytokines and chemokines (IL-1β, IL-6, IL-8, IFN-γ, CCL5, CXCL5, GRO, MCP-1), and tissue repair/regeneration factors (IGF-1, EGF, Leptin, TPO). Compared to 1mln and Matrigel conditions, both EPA and PENTA cultures exhibited significantly elevated secretion of pro-angiogenic and neuroimmune signaling factors. PENTA cultures showed the highest expression of multiple factors, including VEGF-A, IL-6, CXCL5, and IGF-1, suggesting synergistic activation of angiogenic and inflammatory pathways mediated by neuron- and microglia-derived signals. Statistical comparisons reveal that neurovascular complexity is positively correlated with activation of vascular signaling programs. Bars represent mean ± s.e.m. from n = 4-5 independent biological replicates per group. *P < 0.05, **P < 0.01, ***P < 0.001 by one-way ANOVA with Tukey’s post hoc test.

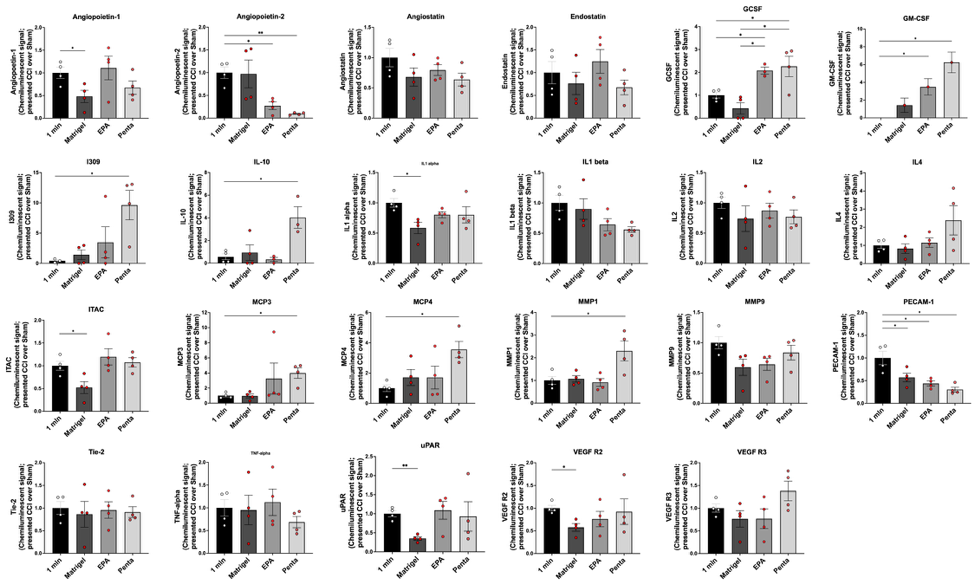

Supplementary Figure 5 - Expansion of angiogenic, extracellular matrix, and inflammatory protein profiles across neurovascular complexity gradients. Bar plots show protein levels of 25 additional markers assessed via multiplex cytokine array in conditioned media from four experimental groups: monoculture brain microvascular endothelial cells (1 mln), Matrigel-embedded BMECs, EPA co-cultures (endothelial cells, pericytes, astrocytes), and fully integrated PENTA cultures (EPA + neurons and microglia). The analytes measured span pro-angiogenic regulators (Angiopoietin-1, -2, -3; VEGF-B, VEGF-C), matrix remodeling enzymes (MMP-1, MMP-8; Enolase-1), neuroimmune markers (GFAP, IL-3, IL-7, IL-15, IL-16, uPAR), and vascular stability signals (PECAM-1, Tie-2, PDGFRβ).

Compared to monocultures and Matrigel-only conditions, EPA and PENTA cultures displayed elevated expression of several matrix remodeling and vascular stabilization markers, including MMP-1, PDGFRβ, and PECAM-1. Notably, PENTA cultures secreted significantly higher levels of Angiopoietin-1, IL-15, and VEGF-C, suggesting that neuron- and microglia-derived signals contribute to the activation of regenerative vascular programs. Conversely, the expression of anti-angiogenic or immune-priming factors such as IL-3 and IL-7 remained low across all groups. Overall, the data demonstrate that increasing neurovascular complexity enhances not only pro-angiogenic cytokine production but also endothelial maturation and vessel-stabilizing signals.

Data represent mean ± s.e.m. for n = 4-5 biological replicates. *P < 0.05, **P < 0.01, ***P < 0.001 by one-way ANOVA with Tukey’s post hoc correction.

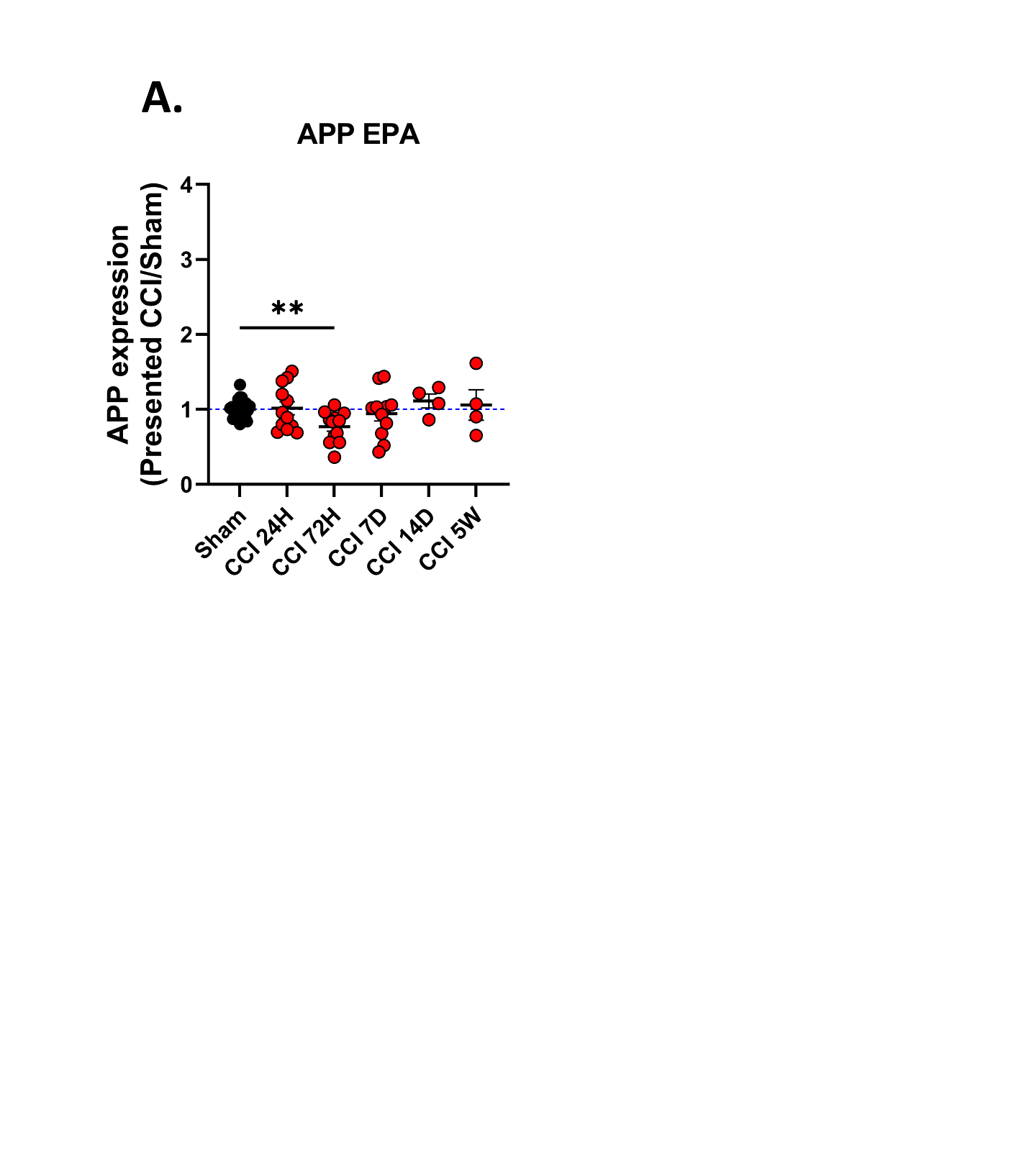

Supplementary Figure 6 – Western blot analysis of EPA moderate injury and sham samples (6 m/s impact, 3mm impact tip) A. Amyloid precursor protein (APP) normalized to total protein content and shown as CCI / sham average.

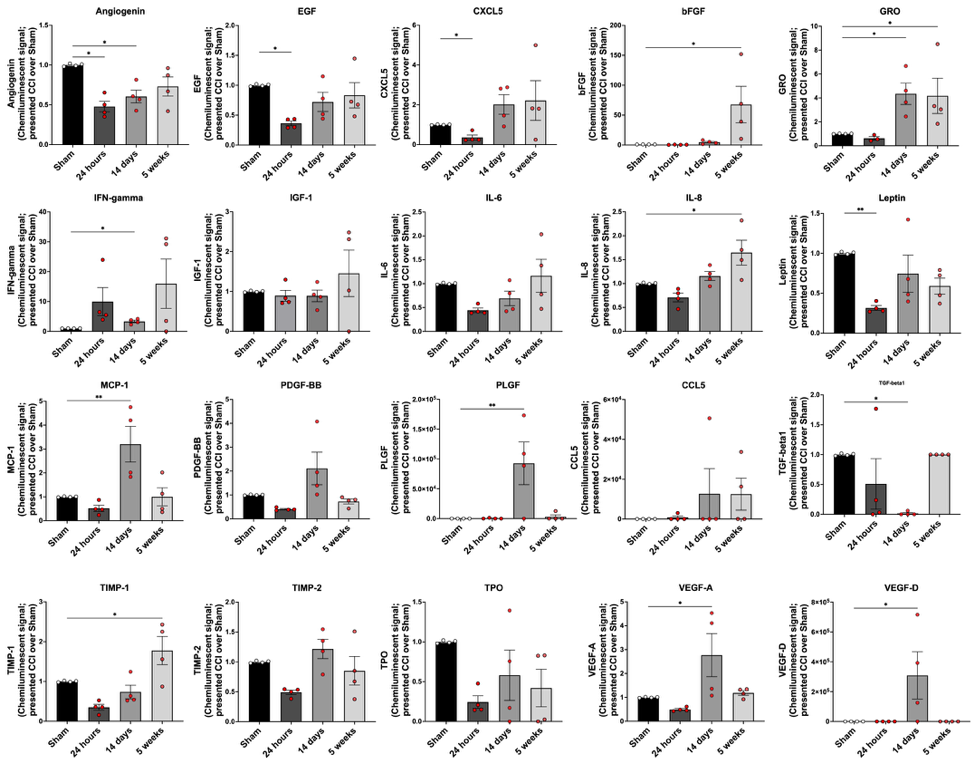

Supplementary Figure 7 - Temporal dynamics of angiogenic and inflammatory mediators in EPA cultures following moderate traumatic injury (Panel 1). Bar graphs show concentrations of 24 secreted proteins quantified from conditioned media of endothelial-pericyte-astrocyte (EPA) cultures at three timepoints following moderate mechanical injury (6 m/s, 3 mm tip, 0.6 mm depth): 24 hours, 14 days, and 5 weeks, compared to sham controls. Analytes include pro-angiogenic growth factors (VEGF-A, VEGF-D, PLGF, bFGF, PDGF-BB), extracellular matrix modulators (TIMP-1, TIMP-2), immune signaling proteins (IL-1β, IL-6, IL-8, IFN-γ, CCL5, CXCL5), and regenerative signals (EGF, IGF-1, Leptin, TPO).

At 24 hours post-injury, several key mediators—including VEGF-A, CXCL5, MCP-1, and IL-6—were significantly elevated, indicating early endothelial activation and inflammatory priming. This pro-inflammatory and pro-angiogenic state partially persisted at 14 days, as evidenced by sustained or increased levels of GRO, IL-8, and IGF-1. By 5 weeks post-injury, some mediators (e.g., bFGF, PLGF) returned to near baseline while others (e.g., GRO, VEGF-D, IL-6) remained elevated, suggesting a chronic low-grade inflammatory state and persistent dysregulation of vascular signaling. Notably, leptin and TPO levels fluctuated non-monotonically over time, reflecting complex metabolic and hematopoietic feedback during injury recovery.

Bars represent mean ± s.e.m. for n = 4-5 biological replicates. *P < 0.05, **P < 0.01, ***P < 0.001 by one-way ANOVA with Tukey’s post hoc test.

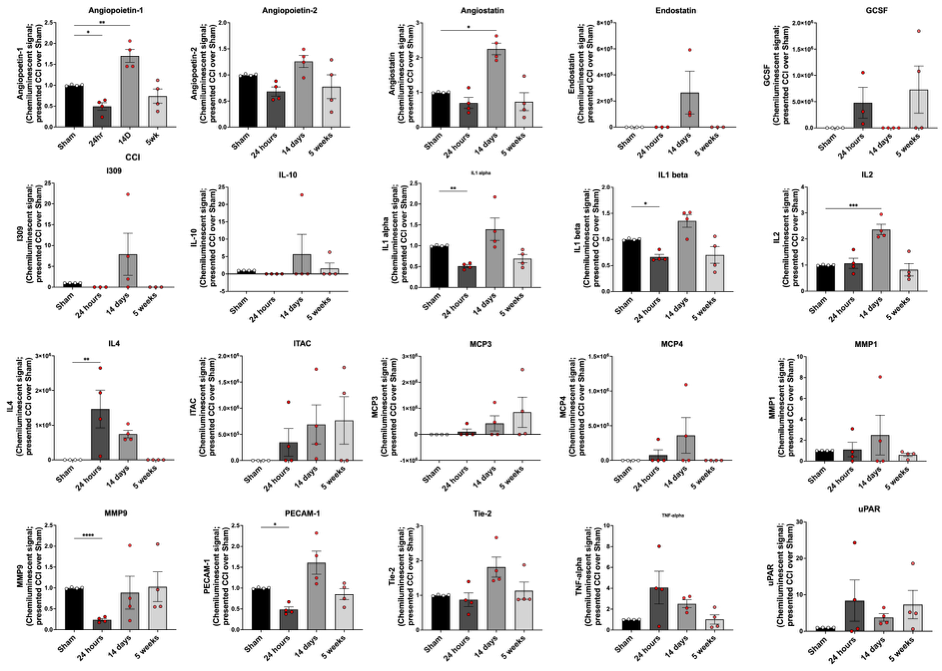

Supplementary Figure 8 - (Panel 2) | Longitudinal profiling of vascular remodeling and inflammatory resolution markers in EPA cultures after traumatic brain injury. Bar plots show concentrations of 25 additional secreted proteins measured from conditioned media of endothelial-pericyte-astrocyte (EPA) co-cultures at 24 hours, 14 days, and 5 weeks after moderate traumatic injury (6 m/s impact velocity, 3 mm tip, 0.6 mm depth), compared to sham controls. This panel focuses on angiopoietins (1-3), endothelial stabilization proteins (Tie-1, PECAM-1), ECM regulators (MMP-1, MMP-8), cytokines (IL-1α, IL-1β, IL-2, IL-3, IL-4, IL-16), growth factors (G-CSF, ENO1), chemokines (MCP-3, MCP-4), and immune response markers (TNF-α, uPAR, IFN-γ).

Angiopoietin-1 and Angiopoietin-2 were consistently upregulated across injury timepoints, while Angiopoietin-3 and PECAM-1 showed delayed increases at 14 days and 5 weeks, indicating sustained or compensatory endothelial remodeling. Pro-inflammatory cytokines including IL-1β, IL-6, and TNF-α peaked acutely at 24 hours and declined by 5 weeks, though levels of IFN-γ and IL-1α remained elevated, suggesting partial immune resolution. Tie-1 expression increased at 14 days and 5 weeks, coinciding with persistent uPAR and MCP-4 secretion. Enolase-1 (ENO1), a marker of metabolic stress and neuronal injury, peaked late, highlighting prolonged metabolic dysfunction in injured cultures.

Overall, this dataset reveals prolonged secretion of endothelial destabilizing factors, chemokines, and inflammatory mediators beyond the acute phase, suggesting that EPA cultures maintain a chronic remodeling state in the absence of neuroimmune regulation. Data are shown as mean ± s.e.m. from n = 4-5 independent replicates. *P < 0.05, **P < 0.01, ***P < 0.001 by one-way ANOVA with Tukey’s post hoc test.

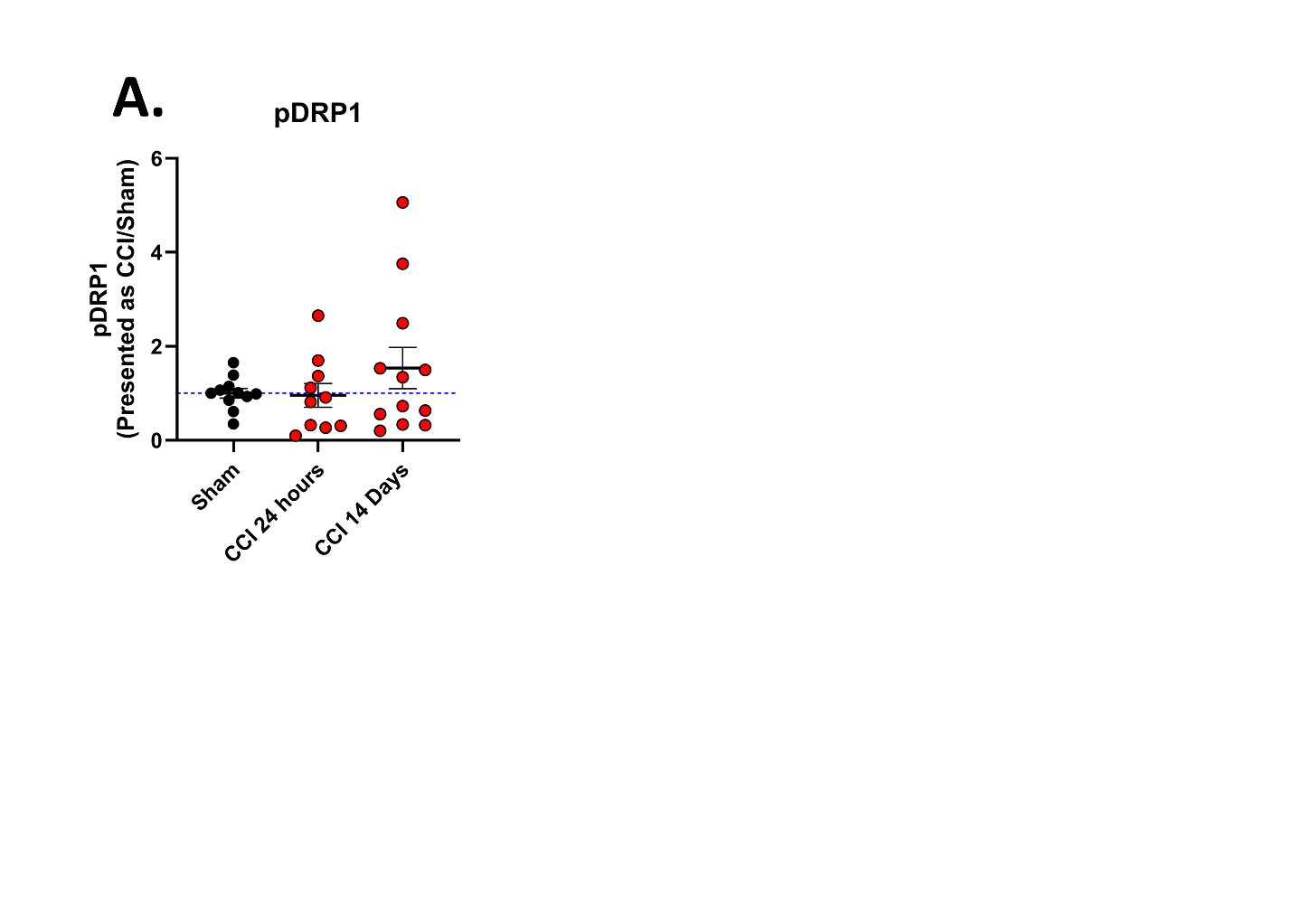

Supplementary Figure 9 – Western blot analysis of PENTA moderate injury and sham samples (6 m/s impact, 3mm impact tip) A. pDRP1 shown as pDRP1 signal divided by DRP1 signal and total protein, normalized by CCI / sham average.

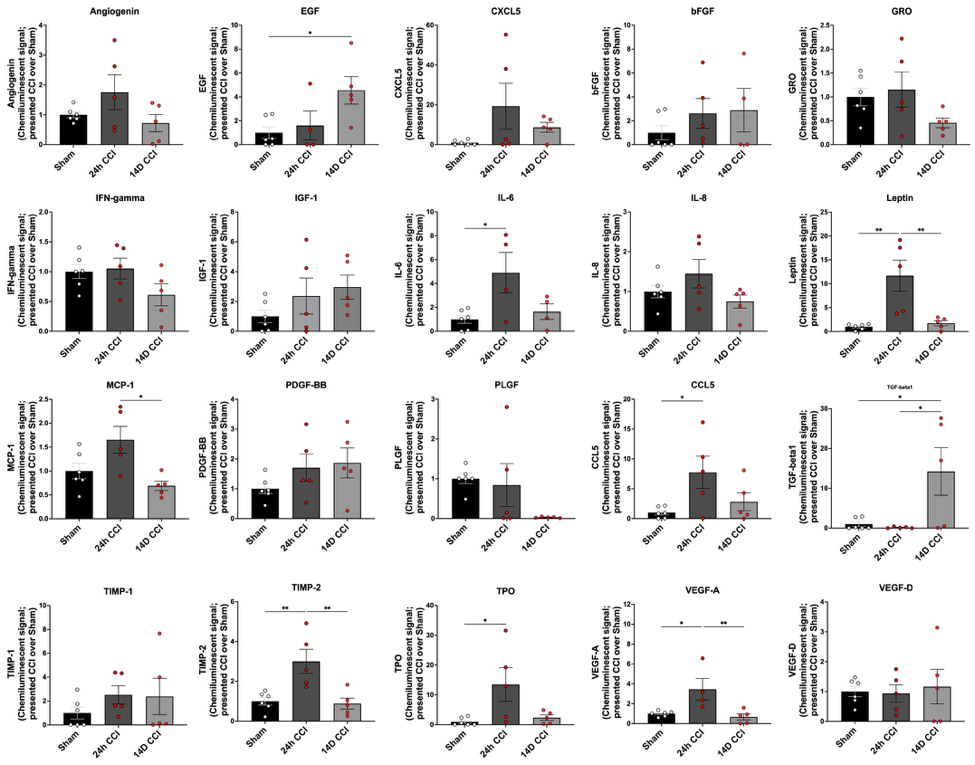

Supplementary Figure 10 – **Individual markers for analysis of angiogenic and immunomodulatory proteins in PENTA cultures following traumatic injury (Panel 1).**
Bar graphs illustrate concentrations of 24 secreted factors in conditioned media from human neurovascular PENTA cultures—comprising endothelial cells, pericytes, astrocytes, neurons, and microglia—measured at 24 hours and 14 days after moderate traumatic brain injury (6 m/s impact velocity, 3 mm tip, 0.6 mm depth), compared to uninjured controls (Sham). Analytes include angiogenic growth factors (VEGF-A, VEGF-D, PLGF, PDGF-BB, bFGF), inflammatory cytokines and chemokines (IL-1β, IL-6, IL-8, IFN-γ, CXCL5, CCL5, GRO, MCP-1), and metabolic and regenerative factors (IGF-1, EGF, Leptin, TPO), as well as matrix remodeling proteins (TIMP-1, TIMP-2).

PENTA cultures exhibited robust acute (24 h) upregulation of multiple angiogenic and inflammatory mediators, including VEGF-A, IL-6, MCP-1, and GRO, consistent with a coordinated vascular injury response. By 14 days, several key mediators remained elevated or were further induced—such as CXCL5, IGF-1, and VEGF-D—highlighting sustained pro-reparative and immune-modulating signaling. Leptin, IL-8, and TPO also showed delayed upregulation, suggesting involvement in longer-term metabolic and hematopoietic adaptations. Compared to EPA cultures, PENTA cultures demonstrated stronger and more temporally coordinated activation of vascular signaling, supporting a role for neuroimmune interactions in amplifying and sustaining post-injury angiogenic remodeling.

Bars indicate mean ± s.e.m. from n = 4-5 biological replicates. *P < 0.05, **P < 0.01, ***P < 0.001 by one-way ANOVA with Tukey’s post hoc correction.

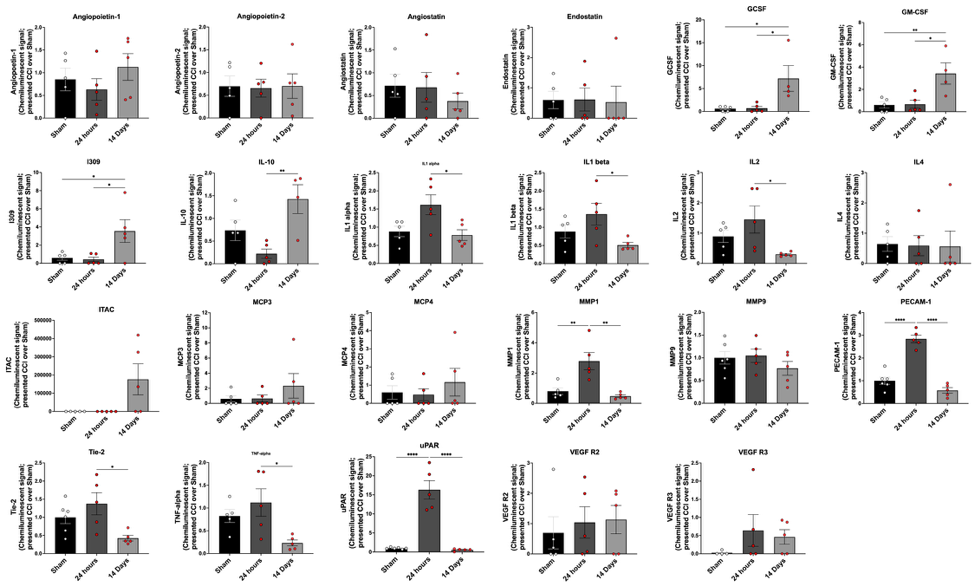

Supplementary Figure 11 - **(Panel 2) | Individual markers for analysis of angiogenic and immunomodulatory proteins in PENTA cultures following traumatic injury.** Bar plots display levels of 24 additional angiogenic, inflammatory, and extracellular matrix remodeling factors measured in conditioned media from PENTA cultures—comprising endothelial cells, pericytes, astrocytes, neurons, and microglia—at 24 hours and 14 days following moderate injury (6 m/s, 3 mm tip, 0.6 mm depth), compared to sham controls. Analytes include angiopoietins (Angiopoietin-1, -2, -3), endothelial stress proteins (Enolase-1), cytokines (IL-1α, IL-1β, IL-2, IL-4, IL-16), colony-stimulating factors (G-CSF, GM-CSF), matrix-associated proteins (MMP-1, MMP-9, PECAM-1), and markers of vascular integrity (Tie-2, VEGF-B, VEGF-C, VEGF-D).

Both acute and sustained elevations in angiopoietin family members and colony-stimulating factors were observed in PENTA cultures, with GM-CSF, G-CSF, and Angiopoietin-2 levels peaking at 14 days post-injury. IL-1 family cytokines (IL-1α, IL-1β), IL-4, and IL-16 also remained elevated, indicating persistent pro-inflammatory signaling. ECM remodeling proteins such as MMP-1 and MMP-9 were upregulated in a time-dependent manner, consistent with dynamic vascular restructuring. Tie-2, VEGF-B, and VEGF-C increased over time, suggesting compensatory activation of endothelial stabilization and vessel maintenance mechanisms. Compared to EPA cultures, the inclusion of neurons and microglia in the PENTA model resulted in more robust and sustained upregulation of immune, angiogenic, and remodeling signals.

Data represent mean ± s.e.m. from n = 4-5 independent biological replicates. *P < 0.05, **P < 0.01, ***P < 0.001 by one-way ANOVA with Tukey’s post hoc test.
